# INSULIN-LIKE GROWTH FACTOR I REDUCES CORONARY ATHEROSCLEROSIS IN PIGS WITH FAMILIAL HYPERCHOLESTEROLEMIA

**DOI:** 10.1101/2022.09.23.509213

**Authors:** Sergiy Sukhanov, Yusuke Higashi, Tadashi Yoshida, Svitlana Danchuk, Mitzi Alfortish, Traci Goodchild, Amy Scarborough, Thomas Sharp, James S. Jenkins, Daniel Garcia, Jan Ivey, Darla L. Tharp, Jeffrey Schumacher, Zach Rozenbaum, Jay K. Kolls, Douglas Bowles, David Lefer, Patrice Delafontaine

## Abstract

**Objective:** Although murine models of coronary atherosclerotic disease (CAD) have been used extensively to determine mechanisms, limited new therapeutic options have emerged. Pigs with familial hypercholesterolemia (FH pigs) develop complex coronary atheromas that are almost identical to human lesions. We reported previously that insulin-like growth factor 1 (IGF-1) reduced aortic atherosclerosis and promoted features of stable plaque in a murine model. We tested IGF-1 effects in atherosclerotic FH pigs to consider use of IGF-1 to treat CAD in humans. FH pigs were administered with IGF-1 for 6 months. Atherosclerosis was quantified by serial intravascular ultrasound (IVUS) and histology, plaque composition - by immunohistochemistry. We used spatial transcriptomics (ST) analysis to identify global transcriptome changes in advanced plaque compartments and to obtain mechanistic insights into IGF-1 effects.

**Results:** IGF-1-injected FH pigs had 1.8-fold increase in total circulating IGF-1 levels compared to control. IGF-1 decreased relative coronary atheroma (IVUS) and lesion cross-sectional area (histology). IGF-1 induced vascular hypertrophy and reduced circulating triglycerides, markers of systemic oxidative stress and pro-atherogenic CXCL12 chemokine levels. IGF-1 increased fibrous cap thickness, and reduced necrotic core size, macrophage content, and cell apoptosis, changes consistent with promotion of a stable plaque phenotype. IGF-1 suppressed FOS/FOSB factors and gene expression of MMP9 and CXCL14 in plaque macrophages, suggesting possible involvement of these molecules in IGF-1’s effect on atherosclerosis.

**Conclusions:** IGF-1 reduced coronary plaque burden and promoted features of stable plaque in a pig model, providing support for consideration of clinical trials. ST profiling of plaques provided novel insights into potential mechanisms.

## INTRODUCTION

Despite lipid lowering and emerging anti-inflammatory agents, atherosclerosis remains the leading cause of death in both men and women in the United States and worldwide^1^. Approximately every 40 seconds, someone in the United States will have a myocardial infarction according to the 2022 heart disease statistics update^2^. The estimated total cost of heart disease in the United States alone is >$329 billion^1^. Thus, interventions that reduce CAD would have substantial health and economic benefits. Coronary atheroma burden is the main determinant of CAD patient outcomes^3^ thus animal models that closely resemble human coronary atherosclerotic disease are of particular interest. Unlike murine models, pigs have plasma lipid profiles very close to humans^4^ and develop coronary disease that is similar to humans^5^. Pigs are the FDA-preferred species for testing human cardiovascular devices and the primary choice for preclinical toxicological testing of anti-atherosclerotic drugs, including statins^6^. Mild atherosclerotic lesions first appear in coronary arteries and both plaque distribution and composition follow a pattern comparable to that of humans^7^, with early lesions transitioning to complex plaques. Familial hypercholesterolemia is a human genetic disorder with high circulating cholesterol and LDL levels, resulting in excessive atherosclerosis^8^. Pigs with familial hypercholesterolemia (FH pigs) have been described by Rapacz and others^9^. FH pigs harbor a point mutation in both LDLR alleles that reduces receptor binding, as well as allele variations in ApoB that may further contribute to the phenotype. FH swine are an excellent model for translational atherosclerosis-related research^10^. Even when consuming a normal diet, FH pigs develop hypercholesterolemia and atherosclerotic lesions ranging from fatty streaks to advanced plaques, with accompanying calcification, neovascularization, hemorrhage, and rupture^7^.

Insulin-like growth factor 1 (IGF-1) is an endocrine and autocrine/paracrine growth factor that has pleiotropic effects on development, metabolism, cell differentiation, and survival. There is ongoing debate on the role of IGF-1 in cardiovascular disease. Traditionally, the role of growth factors in atherosclerosis has been thought to be permissive by stimulating vascular SMC migration and proliferation, thereby promoting neointima formation^11^. Some cross-sectional and prospective studies suggest a positive association between IGF-1 and atherogenesis^12^, but others have found that low IGF-1 is a predictor of ischemic heart disease and mortality, consistent with the potential anti-atherosclerotic, and plaque stabilization effects of IGF-1^13^. Methodological constraints could explain these contradictions because measurement of total IGF-1 levels represents only a crude estimate of the biologically active IGF-1. IGF-1 levels negatively correlate with body weight (BW) and age, and this additionally confounds examination of the role of IGF-1 in CVD.

We have shown that IGF-1 is an anti-atherogenic factor in mice^14^. Systemic IGF-1 infusion reduced aortic root plaque area by ∼ 30% in Apoe^-/-^ mice^15^. This effect is similar to that of statins in the same mouse model (e.g., high-dose rosuvastatin^16^) and statins greatly reduce acute coronary events in patients with atherosclerosis^17^. Consistent with our results, another group reported that long R3 IGF-1 increased plaque smooth muscle cells (SMC), cap thickness and reduced the rate of intraplaque hemorrhage, indicating that IGF-1 promoted plaque stabilization^13^. These findings are in line with most clinical studies^18, 19^ but not all^20^ which have suggested that lower circulating IGF-1 levels and higher levels of IGF-1 binding protein 3 are associated with increased risk of atherosclerotic disease. Confirmation of our murine studies in a large animal model is critical to consider use of IGF-1 to treat atherosclerosis in humans. Anti-atherosclerotic effects of novel drugs are routinely tested in pigs (including statins^21^), however, there are no reports directly testing IGF-1 effects on atherogenesis in a large animal model. Furthermore, since epidemiological studies have reported a positive association between circulating IGF-1 levels and various primary cancers, such as breast, colorectal, and prostate cancer ^22^, it is critical to examine potential carcinogenic effects of long-term IGF-1 administration in a large animal model.

Here we document IGF-1 effects on coronary atherosclerosis in FH pigs, using recombinant human IGF-1 at a dose that is FDA approved for long-term treatment of growth failure in children with severe primary IGF-1 deficiency. Furthermore, we use spatial transcriptomics (ST), an innovative groundbreaking technology that quantifies changes in the whole transcriptome within a morphological context^23^, to detect spatially and differentially expressed genes targeted by IGF-1. Our results provide mechanistic insights into IGF-1-induced effects on atherosclerosis and to our knowledge represent the first report on the use of ST to analyze atherosclerotic tissue from animals or humans.

## METHODS

Extended methods are available in the Supplemental Material. The data, methods, and study materials used to conduct the research will be available from the corresponding author on reasonable request.

### Animals

All animal experiments were performed in accordance with the Guide for the Care and Use of Laboratory Animals, the Public Health Service Policy on the Humane Care and Use of Laboratory Animals, and the Animal Welfare Act. Institutional Animal Care and Use Committee approvals were obtained from the University of Missouri-Columbia and Louisiana State University Health Sciences Center–New Orleans before initiation of experimental studies. The *Rapacz* familial hypercholesterolemic swine (*Sus Scrofa*) (FH pigs) were received from the Swine Research and Teaching Center at University of Wisconsin (Madison, WI). Intravascular ultrasound (IVUS) was performed using the IVUS imaging system (Volcano Corporation, San Diego, CA) and a 20 MHz 3.5 F Visions PV 0.035 Digital IVUS Catheter (Volcano Corporation, San Diego, CA) at baseline (T0) and at 3 months (T3) and 6 months (T6) (Fig.1A). We administered recombinant human IGF-1 (INCRELEX_™_ by IPSEN, 50 ug/kg, SC every 12h, twice per day) to FH pigs for 6 months. The control group received an equal volume of saline. All pigs received a high-fat diet (HFD) starting the day after the T0 IVUS.

**Figure 1.**
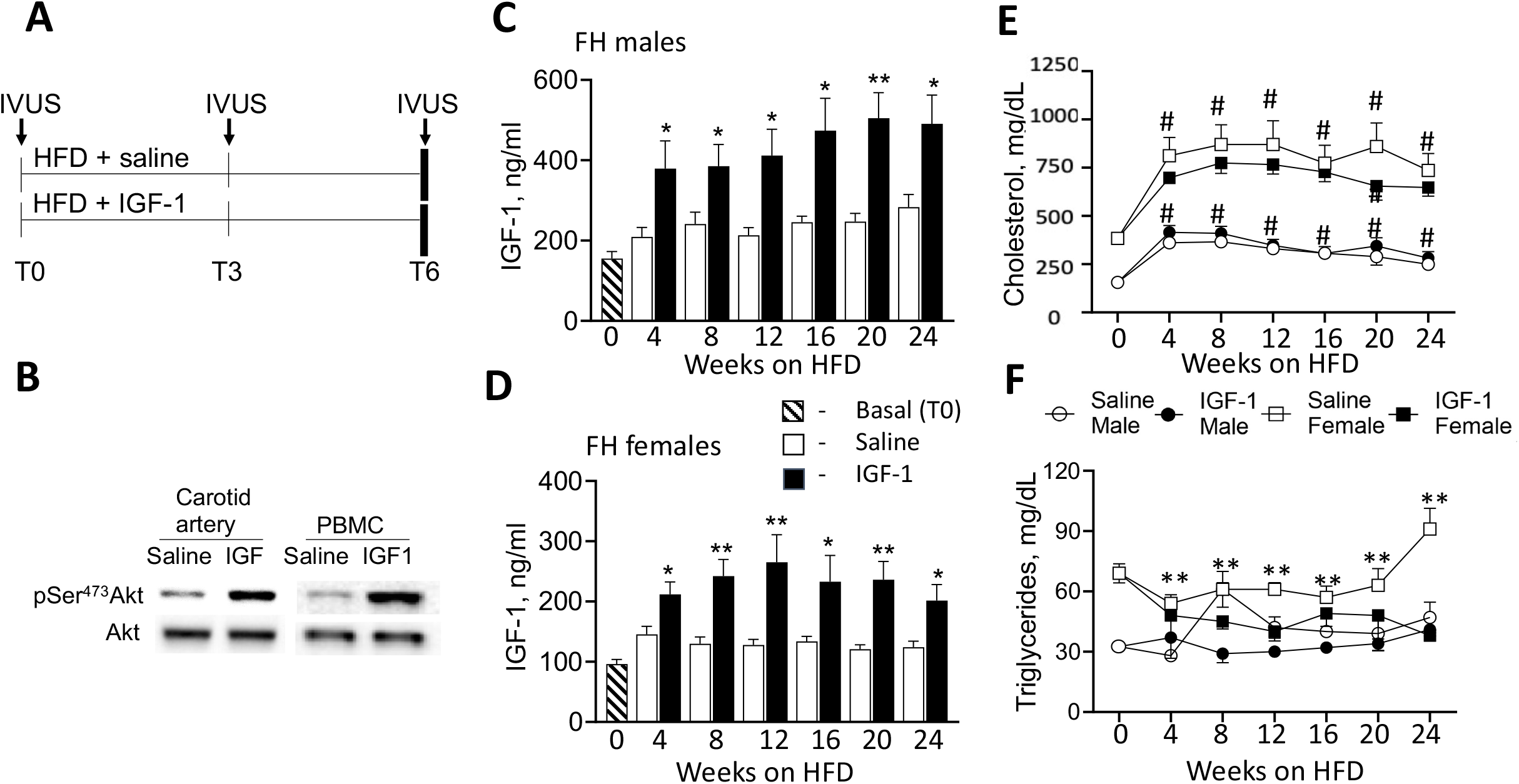
Phenotype of FH pigs. A, Experimental design. FH pigs (males, 5/group; females, 9/group) were injected daily with IGF-1 or saline (control) and fed with high-fat diet (HFD) for 6 months. Coronary atherosclerosis was quantified by intravascular ultrasound (IVUS) before injections (T0), after 3 months (T3) and after 6 months (T6, at sacrificing). B, IGF-1 stimulated specific downstream signaling in porcine carotid artery and in peripheral blood mononuclear cells (PBMC). IGF-1 (or saline) was injected into pig. Tissues and blood were collected 4 hours following injections and Akt phosphorylation was quantified by immunoblotting. C, D, Total plasma IGF-1 level was quantified by ELISA in FH males (C) and females (D). E, F, Cholesterol (E) and triglyceride (F) levels in IGF-1 and saline-injected FH pigs. *P<0.05, **P<0.01 vs. saline, #P<0.05 vs. T0.

### IVUS analysis

For the current study 20 mm of IVUS pullback segment distal to the ostia of the right coronary artery (RCA) and the left anterior descending artery (LAD) were selected. The area circumscribed by the outer border of the echolucent tunica media (TM) and the luminal border was manually traced on each 1 mm IVUS frame within selected fragment. The following indices of vessel morphology were assessed:

1. lumen volume (mm^3^) = lumen area (the area bounded by the luminal border) x length of fragment.
2. vessel volume (mm^3^) = external elastic membrane (EEM) area x length of fragment.
3. plaque + media volume (mm^3^) = vessel volume – lumen volume.
4. relative atheroma volume (%): plaque plus media volume divided by the vessel volume x 100%.

### Atherosclerotic burden and plaque composition analysis

The entire right coronary (RCA) and the left anterior descending (LAD) arteries were collected, and the proximal 30 mm fragment of RCA and LAD was further cut onto 5 mm fragments for embedding in paraffin. Serial 6-μm cross-sections were cut from each histological block and stained with Gomori’s Trichrome stain for morphological analysis. EEM, internal elastic membrane (IEM) and luminal border were manually outlined, and corresponding cross-sectional areas (CSA) were measured. The cellular content of coronary plaques was assessed by immunohistochemistry (IHC) using cell marker-specific antibody for α-SMA, SRA and CD31 to identify smooth muscle-like cells (SMC), macrophage-like cells (MF) and endothelial-like cells (EC), respectively.

### Spatial transcriptomics (ST)

ST utilizes spotted arrays of specialized mRNA-capturing probes containing a spatial barcode unique to that spot. When a cryosection is attached to the slide, the capture probes bind mRNA from the adjacent point in the tissue. After mRNA extraction, cDNA library is generated and sequenced. ST was conducted using the Visium Spatial Gene Expression System (10X Genomics, Pleasanton, CA) in according with manufacturer’s instructions. Pig reference genome was created from *Sus scrofa* genomic sequence (Sscrofa 11.1) and Ensembl annotation, and reads were aligned and counted by Space Ranger (10X Genomics). All the downstream analyses were performed using R toolkit Seurat^24^ and Ingenuity pathway analysis (IPA) software (Qiagen).

### Statistical analyses

Data are presented as mean ± SEM and were analyzed using Excel and GraphPad Prism v.6.0 (GraphPad Software, San Diego, CA). Artwork was generated in GraphPad Prism. Statistical comparisons for histology data were performed by unpaired 2-tailed student’s t test. IVUS results, and blood biochemistry data were analyzed using a three-way repeated measures ANOVA taking treatments (saline vs. IGF-1 administration), time (periodical, repeated measurements), and animal’s sex (female vs. male) as variables. Grubb’s z-test was used to identify outliers. Data sets were first assessed for residuals distribution using D’Agostino-Pearson omnibus normality test and for equal variances using Levene’s test for equality of variances. Differences in outcomes were determined by ANOVA and Bonferroni’s multiple comparisons test, Kruskal-Walls test, or Mann-Whitney U test, accordingly with the normality of residuals distribution. For all comparisons, P<0.05 was considered statistically significant. Adjusted p-value^25^ was used for statistical comparison of genomic data sets obtained with spatial transcriptomics.

## RESULTS

### Phenotype of FH pigs

We used 14-month-old FH non-castrated males (N=5/group) and gilt (never been used for breeding) females (N=9/group) and administered recombinant human IGF-1 (rhIGF-1), 50 ug/kg/day, twice per day, or saline for 6 months (Fig. 1A). Of note, the IGF-1 dose is within the range of the FDA approved dosage for long-term treatment of growth failure in children with primary IGF-1 deficiency^26^ and the amino acid sequence structure of porcine IGF-1 is identical to human IGF-1^27^. IGF-1 levels at T0 were higher in males vs. females (Fig. 1CD, P<0.005). IGF-1 injected pigs have significantly higher IGF-1 levels compared to control pigs at all tested time-points. The average increase in IGF-1 group vs. saline in males was 88.0±19.4% and 83.3±9.7% in females.

Male and female animals had a similar initial body weight (BW) on average and both IGF-1 and saline-injected pigs gained BW steadily throughout the study, with no difference found between saline- and IGF-1 groups (Suppl.Fig.1). FH males had a larger BW increase compared to FH females (M, 69.2±2.9% increase at T6 vs. T0; F, 45.0±5.2% increase, P<0.01).

Blood pressure and heart rate were measured in sedated pigs at each IVUS procedure. We found no difference in systolic/diastolic blood pressure and heart rate between genders, and between saline and IGF-1 groups (Suppl. Table 1). There was no time-dependent change in blood pressure or in heart rate.

To confirm that IGF-1 administration stimulates specific downstream signaling in porcine vasculature and blood cells we injected rhIGF-1 (or saline, control) into a pig and isolated carotid arteries and peripheral blood mononuclear cells (PBMC) after 4 hours. Akt phosphorylation was increased by almost 4-fold (P<0.001) in both vascular tissue and PBMC in IGF-1-injected pigs vs. control (Fig.1B) indicating that IGF-1 promoted specific downstream signaling.

Blood tests were performed monthly. We found no statistically significant difference in complete metabolic profiles and CBC with differential between genders, saline- vs. IGF-1 group (Supplemental Table 2 and 3). However, we found that females had striking 2.3-fold higher cholesterol levels at T0 compared to males (P<0.001, Fig.1E). HFD feeding caused a significant and sustained elevation of total cholesterol levels in both genders. IGF-1 did not change total cholesterol in both males and females. FH females had significantly higher triglycerides levels compared to males at T0 (P <0.005, Fig.1F) and HFD feeding did not change triglycerides. IGF-1 reduced triglycerides in FH pigs (3-way ANOVA, P<0.005, Fig. 1F).

### Necropsy/histopathology findings

IGF-1 levels have been reported to be associated with an increased risk of cancer ^22^, thus we performed autopsies and collected all major organs for histopathological analysis. No gross abnormalities were found either in saline- or IGF-1-injected FH pigs and no tissue was considered carcinogenic by a certified pathologist. All female pigs had multiple coalescing yellow to tan plaques on the intimal surface of the aorta and large visible lipid deposits in right coronary artery (RCA), left anterior descending artery (LAD) and circumflex artery. Two female pigs (saline, 1; IGF-1, 1) had evidence of a myocardial infarction. Abdominal fat deposits, hepatic lipidosis/fatty liver and fatty lymph nodes were found in several saline- and IGF-1-injected females. Two female pigs in the saline group had pleural adhesions and moderate splenic enlargement suggesting inflammation. Sections of the lung, liver, kidney, spleen, lymph node, stomach, small and large intestines, urinary bladder, ovary, uterus, pancreas, salivary gland, skeletal muscle, thyroid, and aorta from each animal were examined. No significant microscopic lesion including inflammatory or neoplastic process was noted in all animals (not shown) except atherosclerosis in the aorta and in coronary arteries.

### IGF-1 decreases coronary atherosclerosis

IVUS was performed in the RCA and LAD at T0, T3 and T6 time-points (Fig.1A). We found no significant difference in vessel volume, lumen volume, and plaque + media volume between RCA and LAD and between genders at T0 (Table 1). There was a time-dependent increase of the vessel volume, presumably due to normal animal growth and to vascular remodeling concomitant with intimal thickening. In fact, the lumen volume time-dependently decreased in both saline- and IGF-1-injected pigs (P<0.001 for both RCA and LAD), consistent with intimal thickening. IGF-1-injected pigs have larger time-dependent increase in RCA and LAD artery volume compared to controls (P<0.05) suggesting vascular hypertrophy. Coronary arteries in IGF-1 injected FH females had larger lumen volume at T3 and T6 (P<0.005) compared to control.

**Table 1.** FH pig coronaries morphology.

| IVUS/drug | Sex/artery | Vessel vol., mm<sup>3</sup> |  | Lumen vol., mm<sup>3</sup> |  | Plaque + media vol., mm<sup>3</sup> |  |
| --- | --- | --- | --- | --- | --- | --- | --- |
|  |  | Mean | SEM | Mean | SEM | Mean | SEM |
| T0 | Males |  |  |  |  |  |  |
|  | RCA | 234 | 14 | 197 | 14 | 36.8 | 1.9 |
|  | LAD | 272 | 13 | 230 | 11 | 41.6 | 3 |
| T3 |  |  |  |  |  |  |  |
| SAL | RCA | 291 | 16 | 224 | 15 | 66.8 | 5.3 |
| IGF-1 | RCA | 320 | 47 | 250 | 38 | 69.1 | 9.5 |
| SAL | LAD | 320 | 5 | 245 | 5 | 75.3 | 3.8 |
| IGF-1 | LAD | 360 | 29 | 285 | 27 | 74.6 | 5.1 |
| T6 |  |  |  |  |  |  |  |
| SAL | RCA | 389 | 28 | 246 | 16 | 146.8 | 12.4 |
| IGF-1 | RCA | 449 | 22 | 308 | 15 | 141.4 | 9.1 |
| SAL | LAD | 469 | 25 | 310 | 15 | 159.5 | 34 |
| IGF-1 | LAD | 438 | 32 | 311 | 24 | 127.6 | 11.9 |
| T0 | Females |  |  |  |  |  |  |
|  | RCA | 192 | 13 | 161 | 12 | 31.1 | 1.9 |
|  | LAD | 231 | 12 | 197 | 10 | 34 | 1.9 |
| T3 |  |  |  |  |  |  |  |
| SAL | RCA | 229 | 20 | 138 | 11 | 91 | 10 |
| IGF-1 | RCA | 258 | 29 | 169 | 22 | 89.1 | 11.2 |
| SAL | LAD | 268 | 12 | 177 | 11 | 91.2 | 9.5 |
| IGF-1 | LAD | 326 | 13 | 232 | 16 | 94.1 | 7.8 |
| T6 |  |  |  |  |  |  |  |
| SAL | RCA | 308 | 31 | 154 | 19 | 154.3 | 13 |
| IGF-1 | RCA | 360 | 31 | 200 | 18 | 160 | 18 |
| SAL | LAD | 349 | 22 | 197 | 13 | 152.8 | 14.2 |
| IGF-1 | LAD | 387 | 18 | 242 | 17 | 144.1 | 12 |

Pigs had approximately 16% of relative atheroma volume at T0. IGF-1 and saline-injected pigs had a time-dependent increase in absolute plaque + media volume and in relative atheroma volume (P<0.001 for RCA and LAD). FH females had significantly larger time-dependent increase in RCA and LAD relative atheroma at T3 (females, 36.6±2.6%, vs. males, 23.3±0.2%) and at T6 (females, 47±4% vs. males, 35.9±1.9%) indicating the presence of a strong gender effect. IGF-1 did not significantly change the absolute plaque + media volume in RCA and LAD. We found no interaction between IGF-1 effect on relative atheroma and gender (3-way ANOVA). As gender does not influence IGF-1’s potential effect on relative atheroma volume, we combined measurements of both sexes and performed a 2-way analysis (Treatment vs. Time) using Bonferroni’s correction for repeated measurements. IGF-1 time-dependently decreased relative atheroma volume in RCA (P<0.05 vs. saline) and in LAD (P=0.054). The IGF-1-induced increase in vessel lumen and reduction in relative atheroma indicate that IGF-1 reduces coronary atherosclerosis.

Trichrome-stained RCA and LAD cross-sections were used for histological analysis. FH females developed larger and more complex plaques in coronaries compared to males in agreement with IVUS results (Fig.3). Plaques in males contained diffuse homogeneous collagen material, lipid droplets, and have neither necrotic cores, nor fibrous caps. We classified plaques in FH males as type III pre-atheroma in accordance with histological classification of human atherosclerotic lesions^28^. Coronary plaques in FH females are significantly larger (RCA: 2-fold increase in cross-sectional area (CSA) vs. males, LAD: 1.5-fold increase, P<0.05, data for saline group), they contain dense fibrous caps, large acellular/necrotic cores, and multiple cholesterol clefts. We observed the presence of strong calcification (Alizarin Red staining, data not shown) and neovascularization (IHC with CD31, endothelial cell marker, data not shown) in coronary plaques in FH females but not in males. Plaques in FH females were classified as advanced type V fibroatheroma^28^.

**Figure 2.**
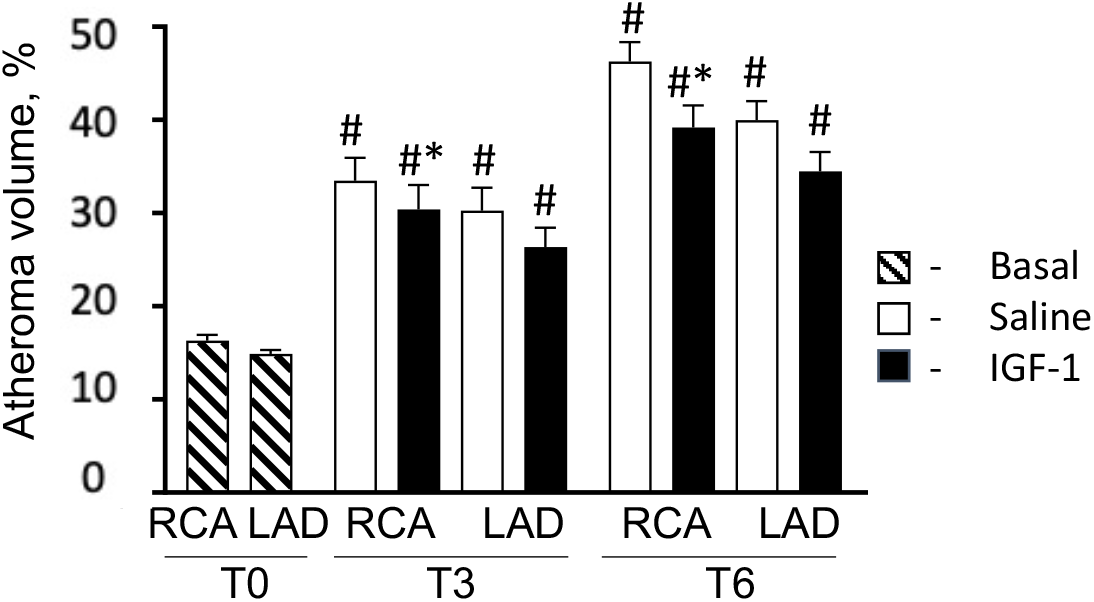
IGF-1 reduced coronary atheroma volume. FH pigs were injected daily with IGF-1 or saline (control) and fed with HFD for 6 months. The coronary atheroma volume was quantified in RCA and LAD by serial IVUS before injections (T0), after 3 months (T3) and after 6 months (T6). RCA and LAD IVUS pullback segments were selected to quantify relative vessel volume, lumen volume and plaque + media volume. Relative atheroma volume (%) was defined as plaque + media volume divided per the vessel volume x 100%. Since gender does not influence IGF-1’s effect on relative atheroma volume, atheroma measurements of both sexes were combined and shown in Figure. *P<0.01 vs. saline, #P<0.05 vs. T0.

**Figure 3.**
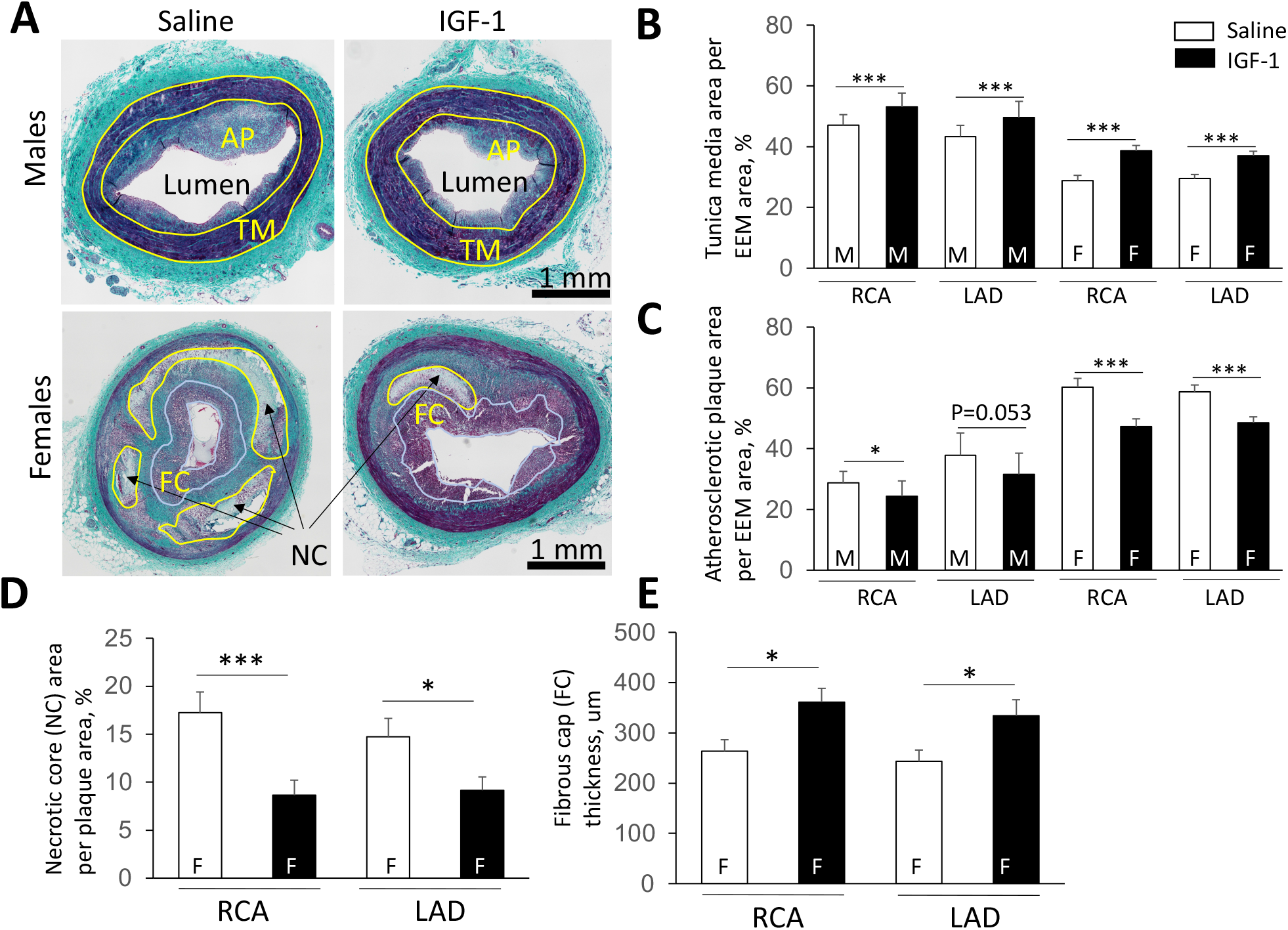
IGF-1 increased vascular media (A, B), reduced atherosclerotic plaque cross-sectional area (C), decreased necrotic core (D) and elevated thickness of fibrous cap (E). RCA and LAD were isolated from IGF-1 and saline-injected FH pigs and coronary artery cross-sections were obtained. A, Representative images of RCA cross-sections obtained from FH males and females. Tunica media (TM), atherosclerotic plaque (AP) and necrotic cores (NC) cross-sectional area were manually outlined on Trichrome-stained sections. The thickness of fibrous cap (FC) was calculated as the mean length of 5 arbitrary lines distributed across the cap area. M, males, F, females, *P<0.05, **P<0.01, ***P<0.005.

FH males have a thicker tunica media compared to females in both IGF-1 and saline-injected groups (P<0.05 in each case, Fig.3B). IGF-1 significantly increased medial CSA (males, RCA: 13% increase vs. control; LAD: 12% increase; females, RCA: 34% increase; LAD: 26% increase) consistent with vascular hypertrophy (Fig.3B). IGF-1 did not change total vessel CSA (outlined by EEM boundary, data not shown). IGF-1 reduced relative atherosclerotic plaque area in males (RCA, 15.2± 4.5% decrease; LAD, 16.6± 7.1% decrease compared to control) and in females (RCA, 21.5±2.7% decrease; LAD, 17.4±2.1% decrease compared to control) (Fig.3C) consistent with IVUS data. Plaques in IGF-1-injected females were more cellular and contained reduced necrotic cores compared to controls (RCA: 50.1±1.6% decrease; LAD: 47.8±1.4% reduction, Fig.3D). IGF-1 significantly increased the thickness of fibrous caps in coronary plaques in female pigs (Fig.3E). Our results indicate that IGF-1 induces vascular hypertrophy, reduces coronary atherosclerotic burden, and promotes features of plaque stability.

### IGF-1 reduces macrophage-like cells and upregulates endothelial-like cells in coronary plaque

We reported previously that IGF-1 increased plaque smooth muscle cells (SMC)^29^, downregulated macrophages (MF) and elevated levels of circulating endothelial cell (EC) progenitors^15^ in HFD-fed Apoe^-/-^ mice. SMC and MF share cell markers in the atherosclerotic plaque^30^ and plaque EC undergo a change in phenotype toward a mesenchymal cell type^31^. Such phenotype switching complicates marker-based cell identification. To validate the IHC protocol, serial RCA sections were stained with a set of cell marker antibodies and immunopositivity pattern was compared. We found that each of 4 SMC marker antibodies stained virtually identical cell population in the plaque and a similar conclusion was made for 3 tested MF and 3 EC marker antibodies (Suppl.Fig.2). These data show that IHC with antibody for a single cell marker identifies plaque cells expressing multiple markers increasing confidence to identify specific plaque cells. We also confirmed that cells immunopositive for SRA, a MF marker were immunonegative for α-SMA, a SMC marker and *vice versa* (Suppl.Fig.2A) showing that these antibodies have no cross-reactivity.

We used α-SMA, SRA and CD31 antibody to quantify SMC-like, MF-like, and EC-like cells, respectively, by immunohistochemistry (IHC). SMC-like cells were abundant in the vascular media and a mixture of SMC- and MF-like cells were found in plaque fibrous cap (Fig.4 and Suppl.Fig.2A). In addition, MF-like cells were present in the area surrounding the plaque necrotic core and colocalized with cholesterol clefts in lipid cores. IGF-1-injected pigs had a slight increase in plaque SMC-like cells (P=NS) (Suppl.Fig.3) and a dramatic 2-fold reduction in plaque MF-like cells in females (P<0.05 for RCA and LAD) (Fig.4). IGF-1 increased EC marker-positive area and this effect reached significance for RCA and LAD in females and LAD in males (Fig.4). Notably, EC marker-positive area was significantly higher in males compared with females (Fig.4A). We noted discontinuous CD31^+^-staining in endothelium layers of both IGF-1 and saline-injected pigs, suggesting either CD31 downregulation, or focal loss of EC, suggesting reduced endothelial integrity. To further obtain a surrogate index of endothelium layer integrity, we normalized CD31^+^ area per lumen perimeter. IGF-1 significantly increased CD31^+^area/lumen perimeter ratio in RCA and LAD in the female group (pixels^2^/pixels, RCA: IGF-1, 5.04±0.84 vs. saline, 2.76±0.64, P<0.05; LAD: IGF-1, 10.31±1.53 vs. saline, 5.52±0.51, P<0.01) suggesting that IGF-1 reduced the number of CD31-positive endothelium layer breaks.

**Figure 4.**
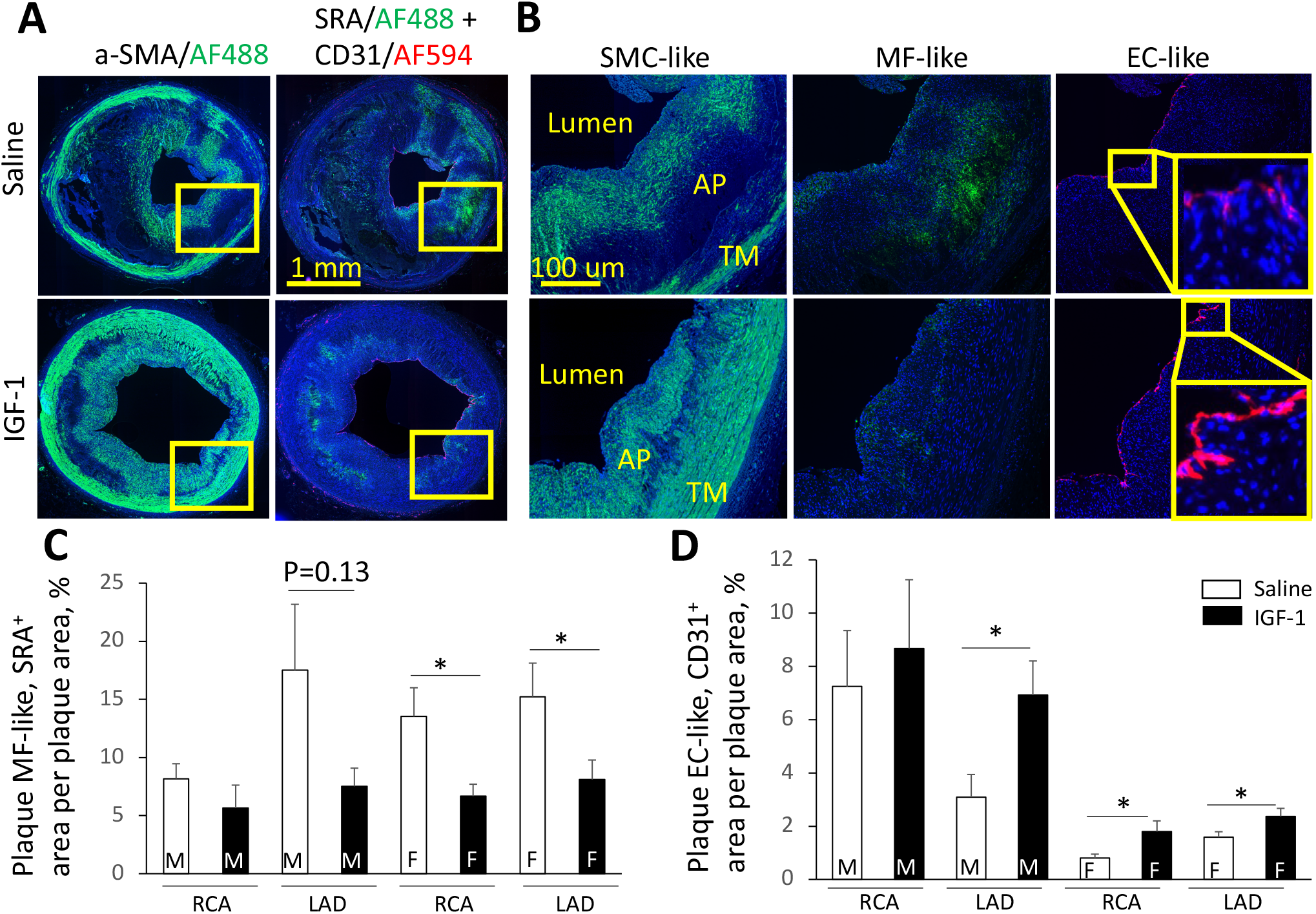
IGF-1 suppressed levels of macrophage-like cells (MF) and upregulated endothelial-like cells (EC) in coronary plaques. Serial RCA and LAD sections were immunostained with α-SMA, SRA and CD31 antibody to identify SMC-like, MF-like and EC-like cells respectively. The primary antibody signal was amplified by biotin/streptavidin or Tyramide systems conjugated to AlexaFluor488 (for α-SMA and SRA) or AlexaFluor594 (CD31). A, representative images of RCA sections obtained from IGF-1 or saline-injected FH females. Yellow square in panel A outlines plaque area magnified in panel B. AP, atherosclerotic plaque, TM, tunica media. B, SMC-, MF- and EC-marker immunopositive cells. C, D, quantitative data. M, males, F, females, *P<0.05 vs. saline.

Systemic IGF-1 administration increased expression of pro-α1(I) collagen in aortic lysates^32^ and SMC-specific IGF-1-overexpression increased collagen fibrillogenesis in the atherosclerotic plaque in Apoe^-/-^ mice^33^. We found that IGF-1 injected pigs had a trend toward increased collagen levels in the vascular media and in coronary plaques (approx.10% increase, P=NS) (Suppl.Fig.3BC). Thus, IGF-1 changed the cellular composition of porcine coronary plaques. Atherosclerotic lesions in IGF-1-injected pigs had decreased levels of MF-like cells and increased endothelial-like cells compared to controls.

### IGF-1 decreases cell apoptosis, reduces systemic oxidative stress, and suppresses inflammation

IGF-1 is a mitogen and pro-survival molecule^34^ and exerted anti-apoptotic, antioxidant and anti-inflammatory effects in Apoe^-/-^ mice^15, 29^. Apoptotic cells in porcine coronary plaques were localized on the luminal border, in the fibrous cap and around necrotic cores (see insert in Fig.5A). IGF-1-injected pigs had an almost 3-fold decrease in cell apoptosis in the male group (P<0.05 for LAD) and approximately 2-fold reduction in apoptosis rate in females (Fig.5D). Proliferating cell nuclear antigen (PCNA) is a cell proliferation marker. Atherosclerotic plaques in the female group had increased PCNA-immunopositivity compared to males (Fig.5E). IGF-1 upregulated PCNA levels in coronary plaques in females (P<0.05 in RCA) and did not change PCNA signal in the male group.

**Figure 5.**
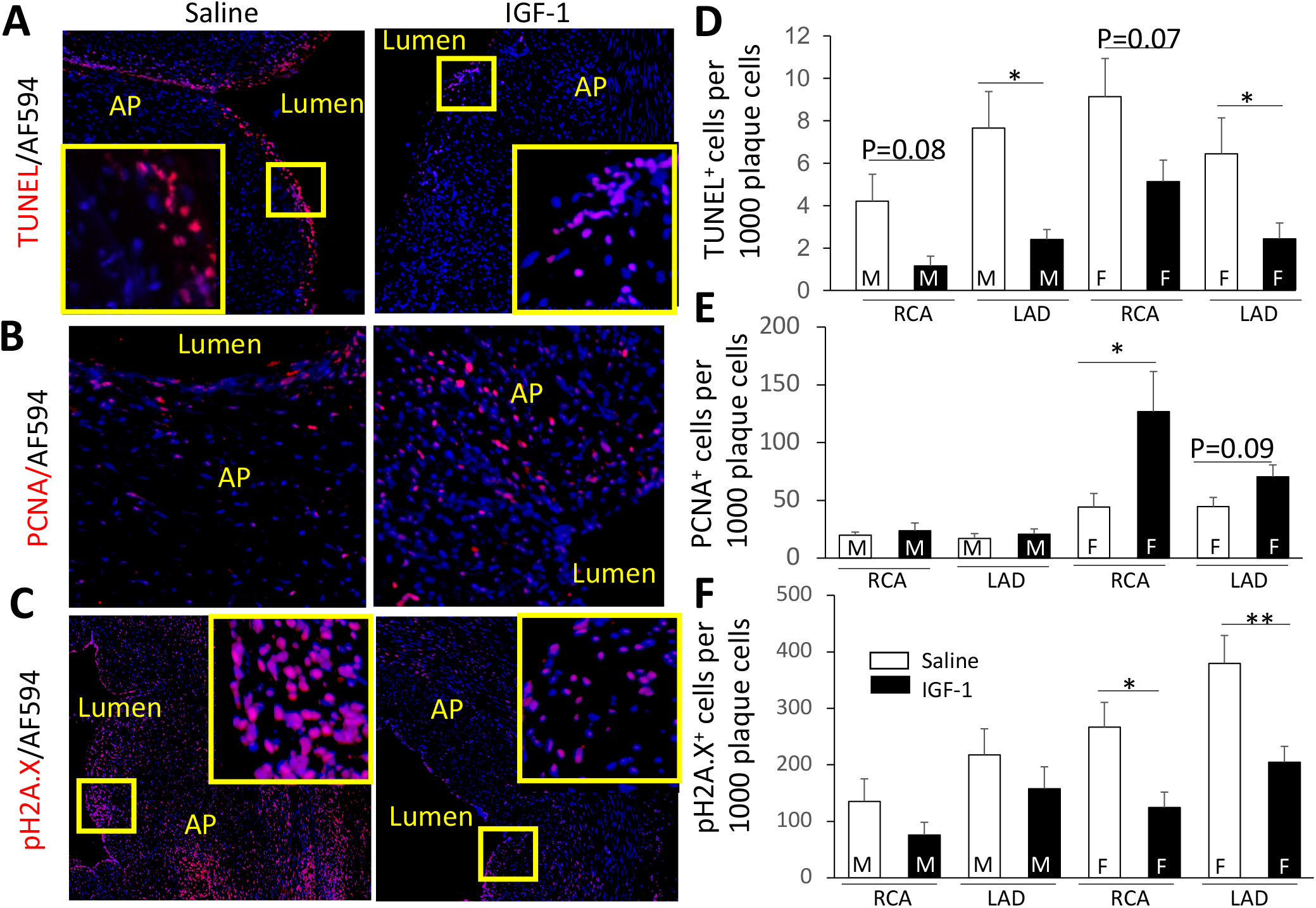
IGF-1 reduces cell apoptosis (A, D), increases cell proliferation (B, E) and suppresses DNA damage (C, F). Cell apoptosis was quantified by TUNEL assay, cell proliferation - by immunostaining with PCNA antibody, and DNA damage - by immunohistochemistry with pH2A.X antibody. A, B, C, representative images of RCA sections obtained from IGF-1 or saline-injected FH females. D, E, F, quantitative data. M, males, F, females, *P<0.05 vs. saline.

Oxidative stress is a major characteristic of hypercholesterolemia-induced atherosclerosis^35^. Oxidative DNA damage promotes cell apoptosis and contributes to formation of unstable plaques. Histone H2A.X phosphorylation is a highly specific molecular marker to quantify DNA damage^36^. We found that 15-35% of plaque cells in coronaries contained detectable levels of pH2A.X (Fig.5C) and higher pH2A.X levels correlated with larger atherosclerotic burden seen in females. IGF-1 significantly decreased the number of pH2A.X^+^ cells in plaques in the female group. Circulating levels of N-tyrosine and total plasma antioxidant capacity (TAC) were measured as indices of systemic oxidative stress (Fig.6). IGF-1 decreased plasma N-tyrosine levels in females at both T3 and T6 time-points (56% and 47% decrease, respectively vs. saline, P<0.05) (Fig.6A). FH female pigs had a 2-fold reduction in TAC compared to males at T6. IGF-1 upregulated TAC in males and females (Fig.6B), although the increase in males did not reach statistical significance. Taken together the N-tyrosine and TAC data indicate that IGF-1 suppressed systemic oxidative stress.

**Figure 6.**
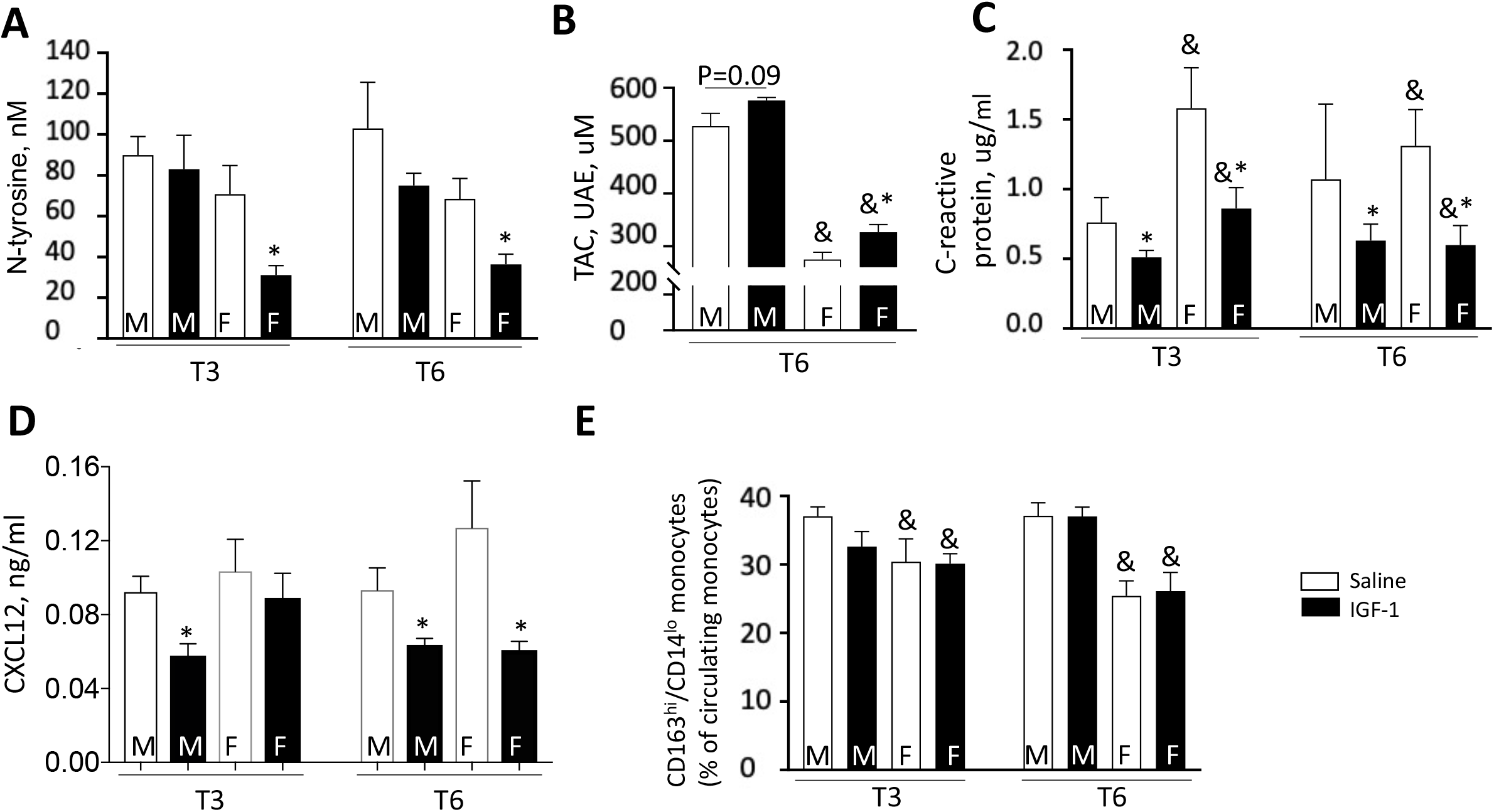
IGF-1 downregulates markers of systemic oxidative stress (A, B), decreases C-reactive protein (C) and chemokine CXCL12 (D). IGF-1 did not alter circulating monocytes subsets (E). Circulating N-tyrosine, CXCL12 and C-reactive protein levels were quantified by ELISA and total antioxidant capacity (TAC) - by using colorimetric assay. TAC assay results shown in urinary acid (standard) equivalents (UAE). E, Whole blood was mixed with a cocktail of antibodies against CD163-PE, CD14-Alexa Fluor 488, and porcine CD172a and subsequently with streptavidin-APC/Cy7. CD172a-positive leukocytes were size-gated and further differentiated into subsets based on CD163 and CD14 expression levels by using FACS. M, males, F, females, *P<0.05 vs. saline, ^&^P<0.05 vs. males.

C-reactive protein (CRP) is an acute marker of inflammatory responses and circulating levels of CRP correlate with progression of CAD^37^. Human CRP transgene expression caused accelerated aortic atherosclerosis in Apoe^-/-^ mice^38^. FH females had higher circulating CRP levels compared to males. IGF-1 significantly decreased CRP levels of both sexes at T3 and T6 (Fig.6C) suggesting a reduction of inflammatory responses. Macrophage-specific IGF-1 reduced chemokine CXCL12 levels, and this effect was associated with decreased atherosclerotic burden in Apoe^-/-^ mice^39^. IGF-1 significantly reduced circulating CXCL12 in FH males at T3 and T6 time-points and in female group at T6 (Fig.6D).

The frequency of monocyte subsets has been linked to severity of atherosclerosis in patients with stable coronary heart disease^40^. We measured two subsets of circulating monocytes, defined by surface expression levels of CD163 and CD14. Monocyte subpopulations were assessed by flow-cytometry. CD172a-positive myeloid cells were size-gated to monocytes, which were further assessed for CD163 and CD14 expression levels, classifying cells to CD163^hi^/CD14^lo^ monocytes and CD163^lo^/CD14^hi^ monocytes. The population size of CD163^hi^/CD14^lo^ monocytes was larger in male animals than in female animals (P<0.001) (Fig.6). No significant effects were induced by IGF-1 administration.

### IGF-1 changes the global transcriptomic profile of coronary plaques

To obtain further insights into the effects of IGF-1 on coronary atherosclerosis we performed an exploratory analysis of advanced plaques from FH females, using the newly developed technology, spatial transcriptomics (ST). ST provides an unbiased picture of entire transcriptome changes within a spatial context^23^. Prior to running ST analyses we confirmed the quality of RNA preparations. Total RNA was extracted from plaque-containing RCA cryosections (N=4), and RNA integrity number (RIN)^41^ was quantified using an Agilent Bioanalyzer. RIN was >7, considered suitable for ST analysis according to the manufacturer’s (Visium/10X genomics) recommendations. RCA cryosections from IGF-1- and saline-injected FH females (N=2/group) were processed and ST quality controls (Suppl.Table 4) were consistent with a successful experiment. Correlation of ST gene expression with protein expression (measured by IHC) was confirmed for selected gene/protein combinations including IGF-1 binding protein 7 and α-SMA (data not shown). Furthermore, changes in gene expression of MMP9, IGFBP7, and FOS measured by ST were confirmed by RT-PCR using aliquots of mRNA isolated from tissue sections (data not shown).

We performed unsupervised clustering of all ST spots in IGF-1 and saline specimens. ST spots were grouped into 9 clusters (0-8) based on their transcriptome. We identified the top 10 genes overexpressed in each cluster (vs. all other clusters) to obtain the heatmap (Fig. 7A). In parallel, plaque fibrous cap (FC), lipid core, tunica media and tunica adventitia were outlined using H&E images and transcriptome clusters and histological annotations were compared side-by-side (Fig.7 A, B, Suppl. Fig.4). Fibrous cap (FC) contained mainly (94.4%) cluster 1 and 2, and tunica adventitia cluster 0 and 3 (Suppl.Fig.4A).

**Figure 7.**
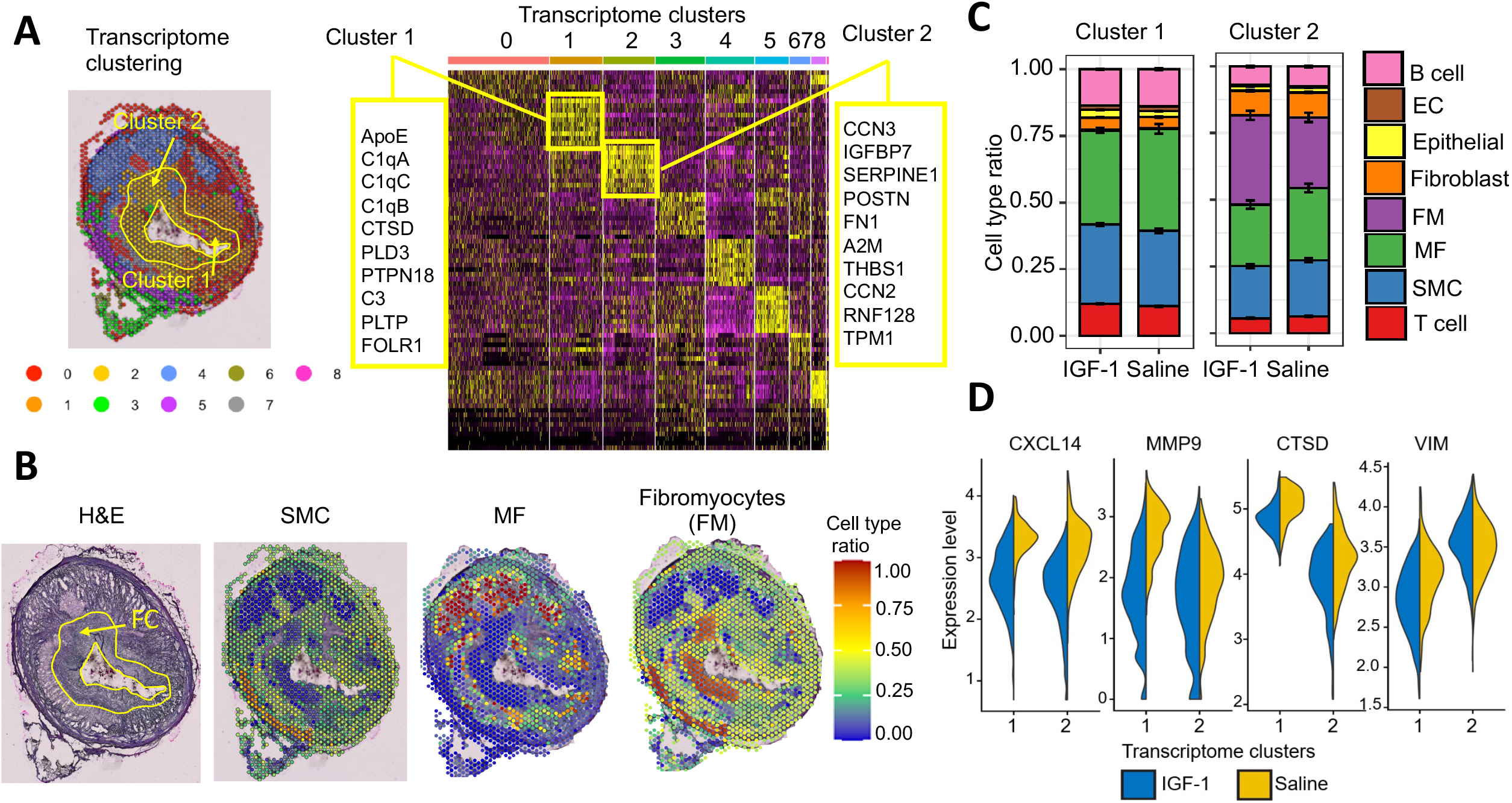
IGF-1 alters global transcriptomic profile of coronary plaque. The RCA cryosections were obtained from IGF-1- (N=2) and saline-injected (N=2) FH females and spatially resolved global transcriptome was assessed by spatial transcriptomics (ST). A, ST spots in IGF-1 and saline specimens were grouped into 9 clusters (0-8) based on their transcriptome and the heat map with top 10 upregulated genes/cluster was generated. B, Cell type ratio was calculated for each ST spot using mixed cell deconvolution algorithm to identify spots enriched by SMC, MF or fibromyocytes (FM). C, Cluster 1 and 2 were detected in fibrous cap (FC) and cell type ratio was calculated for IGF-1 vs. saline specimens. D, Violin plots for expression levels of CXCL14, MMP9, CTSD (cathepsin D) and VIM (vimentin) comparing IGF-1 vs. saline specimens in cluster 1 and 2. MMP9, CTSD and VIM were among the top differentially expressed genes between cluster 1 and 2 and were selected based on their atherosclerosis relevance. IGF-1 downregulated CXCL14, MMP9, VIM and CTSD within cluster 1 and decreased expression of CXCL14, MMP9, and CTSD within cluster 2 compared to saline.

The mixed cell deconvolution algorithms use single cell RNA sequencing (scRNAseq) data as a reference to characterize cellular heterogeneity in spatial context^42^. However, to our knowledge there is no scRNAseq data available for pig atherosclerotic specimens. Therefore, we used scRNAseq data obtained for human atherosclerotic RCA (GSE131778^43^) as a reference to assess the cellular composition in our porcine ST dataset. Using deconvolution algorithm, we calculated cell type ratio for each ST spot to identify spots enriched in SMC (SMC-high spots), MF (MF-high) or fibroblast-like cells, termed fibromyocytes (FM-high) (Fig.7B). Of note, we observed good agreement between localization of ST-defined SMC- and MF-high spots and IHC-detected immunopositivity pattern obtained for cell markers. The cell type ratio shows that transcriptomic cluster 4 contains around 80% MF-high spots, cluster 5 has >70% SMC-high spots and virtually all FM-high spots (>95%) were assigned to cluster 2 (Suppl.Fig.4B).

Table 2 shows the top up- and down-regulated genes in IGF-1 vs. saline specimens identified in all ST spots and Table 3 contains a list of differentially expressed genes in SMC-, MF- and FM-high spots. IGF-1 dramatically (>10-fold reduction vs. saline) downregulated FOS and FOSB protooncogenes in all ST spots and in SMC-high spots. IGF-1 downregulated expression of the cytoskeletal molecule desmin in all spots (Table 2) and upregulated ribosomal protein S27 (Table 2), a component of the translational machinery^44^. Activation of translation is in line with known growth-stimulating IGF-1 action^34^. Metalloproteinase 9 (MMP9) gene expression level was reduced by IGF-1 in all ST spots and in MF-high spots. This is consistent with our recent report that MF-specific IGF-1 overexpression downregulated MMP9 in peritoneal MF and decreased aortic atherosclerosis in Apoe^-/-^ mice^45^. Cytokine CXCL14 was the top IGF-1 downregulated gene in MF-high spots (Table 3). CXCL14 was upregulated in MF-derived foam cells and CXCL14 peptide-induced immunotherapy suppressed atherosclerosis in Apoe^-/-^ mice^46^ suggesting a pro-atherogenic role of CXCL14.

**Table 2.**
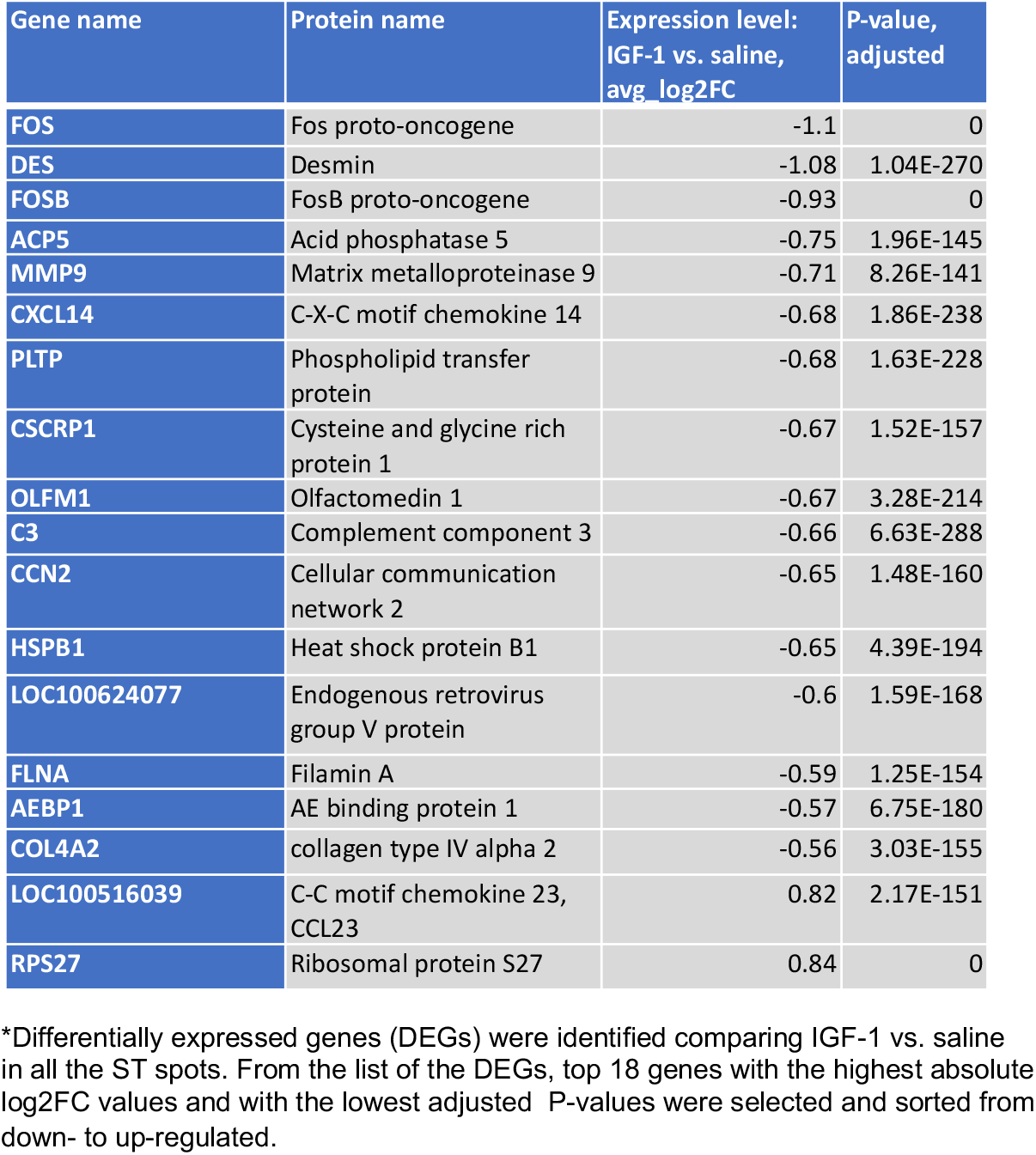
IGF-1 vs. saline top differentially expressed genes in all spatial transcriptomic (ST) spots*.

**Table 3.** IGF-1 vs. saline top differentially expressed genes in cell-high spatial transcriptomic (ST) spots**.

|  | Gene name | Protein name | Expression level<br>IGF-1 vs. saline,<br>avg_log2FC | P-value,<br>adjusted |
| --- | --- | --- | --- | --- |
| SMC-high spots | FOSB | FosB proto-oncogene | -1.85 | 1.36E-20 |
|  | FOS | Fos proto-oncogene | -1.82 | 3.66E-21 |
|  | CRYAB | Crystallin alpha B | -1.05 | 1.57E-08 |
|  | LMNA | Lamin A/C | -0.95 | 1.22E-07 |
|  | HSPB1 | Heat shock protein family B (small) member 1 | -0.94 | 3.99E-16 |
|  | FLNA | Filamin A | -0.85 | 9.04E-14 |
|  | CNN1 | Calponin 1 | -0.67 | 7.50E-05 |
|  | AEBP1 | AE binding protein 1 | -0.58 | 2.10E-07 |
|  | MGP | Matrix Gla protein | 1.06 | 3.23E-19 |
|  | S100A1 | S100 calcium binding protein A1 | 1.11 | 9.54E-07 |
| MF-high spots | CXCL14 | C-X-C motif chemokine ligand 14 | -0.87 | 3.44E-16 |
|  | APOE | Apolipoprotein E | -0.72 | 6.70E-26 |
|  | DDIT4 | DNA damage inducible transcript 4 | -0.59 | 3.01E-26 |
|  | MMP9 | Matrix metalloproteinase 9 | -0.51 | 1.17E-18 |
|  | SIRPA | Signal regulatory protein alpha | -0.51 | 3.15E-12 |
|  | ID2 | Inhibitor of DNA binding 2 | 0.54 | 5.95E-20 |
|  | RPS15A | Ribosomal protein S15A | 0.63 | 3.69E-17 |
|  | CTSL | Cathepsin L | 0.79 | 5.65E-16 |
|  | LGALS3 | Galectin 3 | 0.91 | 2.27E-14 |
|  | RNF128 | Ring finger protein 128 | 1.10 | 8.56E-12 |
| FM-high spots | MGP | Matrix Gla protein | 0.58 | 4.07E-10 |
|  | GSN | Gelsolin | 0.86 | 0.008 |
|  | PENK | Proenkephalin | 1.20 | 0.017 |
|  | S100A2 | S100 calcium binding protein A2 | 1.27 | 0.024 |

\*\*SMC-, MF- and FM-high ST spots were identified by mixed cell deconvolution algorithm driven by Seurat and using human atherosclerotic RCA single cell RNA sequencing data-set as a reference. Top genes were selected by the lowest adjusted P-value and sorted from down- to up-regulated.

We observed a clear boundary between cluster 1 and 2 within the same histological area (FC, Fig.7A), showing the capability of ST analysis to identify plaque areas with different gene expression patterns, that are not histologically distinguishable. To take advantage of this unique ST feature and considering the importance of the FC for overall stability of atheroma we focused our subsequent analysis on cluster 1 and 2. First, we found discrete differences in the cellular composition of cluster 1 and 2. There are more T cells, B cells and MF in cluster 1, whereas fibroblasts levels were lower (Suppl.Fig.4B). The top upregulated genes in cluster 1 (vs. other clusters) includes components of the complement system (C3, C1qA, C1qC and C1qB), and cathepsin D (Fig. 7A). The top genes in cluster 2 includes CCN2 and CCN3 (CCN molecules are involved in wound healing and fibrosis^47^) and fibronectin 1, which is known to play a vital role in tissue repair^48^. A comparison of differentially expressed genes between cluster 1 and 2 showed that upregulated cathepsin D (CTSD) was the top-ranked differentially expressed gene in cluster 1 vs. 2 (∼2.1-fold higher expression in cluster 1 vs. 2; adjusted p-value=3.5×10^−225^). Cathepsin D is a pro-apoptotic molecule and collagen-degrading protease that is highly expressed in MF^49^. Higher cathepsin activity is associated with unstable plaque^49,^. Ingenuity Pathway Analysis (IPA, QIAGEN) predicted that necrosis and apoptosis pathways are upregulated in cluster 1 vs. 2 (data not shown) consistent with increased expression of cathepsin D. Importantly, cluster 1 is the thinnest part of the FC. Taken together these data suggest that cluster 1 is a tissue-degrading and less-fibrotic compartment of the FC compared to cluster 2. We speculate that cluster 1 represents a plaque site with potentially increased vulnerability and propensity to erode or rupture.

We also compared cell type ratio and DEGs in cluster 1 and 2 in IGF-1 vs. saline specimens. IGF-1 slightly decreased MF in both cluster 1 and 2 and IGF-1 upregulated FM in cluster 2 (Fig.7C). SMC transition into FM prevents FC thinning^43^ suggesting that higher FM levels in the FC are beneficial during the disease process. IGF-1 reduced cathepsin D expression in both clusters, whereas vimentin expression (VIM) was decreased only in cluster 1. The top-ranked DEGs in comparing IGF-1 vs. saline groups in cluster 1 and 2 included downregulated chemokine CXCL14, and MMP9 (Fig. 7D). Thus, use of ST combined with deconvolution algorithms provided, to our knowledge for the first time, a profile of the spatial transcriptome of advanced atherosclerotic plaque, and changes induced by IGF-1.

## DISCUSSION

To our knowledge the current study is the first to test the effects of IGF-1 given long term in a large animal model of atherosclerosis. We found that IGF-1 induced vascular hypertrophy, reduced coronary atheroma volume, and decreased plaque cross-sectional area over 6 months and that there was no evidence of IGF-1 induced tumorigenesis over this period. There was evidence of significant sex-specific differences in atherosclerosis development. Females had higher cholesterol and triglyceride levels and an advanced plaque phenotype. IGF-1 increased fibrous cap thickness, and reduced necrotic core size, macrophage content, and cell apoptosis, changes consistent with promotion of a more stable plaque phenotype. IGF-1 also reduced circulating triglycerides, and markers of systemic oxidative stress. ST analysis of plaques from female FH pigs, combined with mixed cell deconvolution algorithm, allowed us to spatially profile gene expression in advanced coronary plaques, and to identify differentially expressed genes in IGF-1 treated animals. We detected nine specific gene clusters and found that the plaque fibrous cap was comprised almost exclusively of cluster 1 and 2, showing the unique capability of ST to identify plaque sub-compartments that are not histologically distinguishable. IGF-1 induced marked decreases in FOS/FOSB transcription factor and in MMP9 and CXCL14 gene expression in plaque macrophages, suggesting involvement of these molecules in IGF-1 anti-atherogenic effects.

Although multiples studies have shown that IGF-1 exerts anti-atherosclerotic effects in murine models^13, 14^ the role of IGF-1 in the development of human atherosclerotic disease is unclear^18, 19^. Both IGF-1 administration and gain-of-function and loss-of-function approaches in mice have provided mechanistic insights, but consideration of IGF-1 for treatment of cardiovascular disease in humans mandates demonstration of IGF-1 efficacy and safety in a large animal model that is physiologically closer to humans.

We administered IGF-1 at a dose approved for use in children with primary IGF-1 deficiency^26^. FH males were non-castrated to fully evaluate potential sex-dependent effects. Coronary plaque location, size and frequency (as assessed by coronary angiography, data not shown) were similar to ones reported for humans^50^. Coronary atheroma cellular composition, the presence of large necrotic/lipid cores, neovascularization, and calcification in FH females closely resembles the phenotype of advanced fibroatheromas reported for patients with coronary atherosclerotic disease (CAD)^51^. These data demonstrate that FH pigs are a valuable model to study CAD and test anti-atherosclerotic drugs.

The number of pre-clinical studies comparing plaque development between the sexes is extremely limited relative to the vast literature exploring atherosclerosis mechanisms^52^. FH females had reduced circulating IGF-1 levels, and higher plasma cholesterol and triglyceride levels compared to males. HFD feeding caused a significant and sustained elevation of total cholesterol levels in both genders. The causative role of high cholesterol in promoting atherogenesis has been well established using multiple animal models including miniature pigs^53, 54^ and genetically modified mice and rabbits^55, 56^. High cholesterol and triglycerides are classical risk factors for human CAD^57^. We hypothesize that high lipid levels in FH females were a major cause of the increased atherosclerosis in this group, although it is possible that lower basal IGF-1 levels in female pigs also increased susceptibility to atherosclerosis development. Of note, although IGF-1 had no effect on total cholesterol it reduced triglycerides (Fig. 1F).

We demonstrated that IGF-1 exerted atheroprotective effects on pre-atheroma (in males) and advanced fibroatheromas (in females). Accumulating evidence suggests that endothelial dysfunction is an early marker for atherosclerosis^58^ and damage of coronary endothelium has been shown to constitute an independent predictor of cardiovascular events^58^. The barrier function of the endothelium is impaired in atherosclerosis leading to uncontrolled leukocyte extravasation and vascular leakage^59^. Intriguingly, we found that the IGF-1-induced anti-atherosclerotic effect in FH pigs was associated with reduced EC damage. Indeed, we found multiple breaks in the EC layer in porcine plaque indicating loss of endothelial integrity. IGF-1-injected pigs had a reduced number of EC layer breaks in coronary plaques suggesting improved EC function. This result is in line with our recent report showing that EC-specific IGF-1 receptor deficiency downregulated endothelial intercellular junction proteins, elevated endothelial permeability and enhanced atherosclerotic burden in Apoe^-/-^ mice^60^. Increased EC lining in plaques in IGF-1-treated is potentially due to elevated EC proliferation or increased recruitment of circulating EC progenitors to the plaque area. IGF-1 has been shown to promote EC proliferation^61^, and increase levels of EC progenitors in Apoe^-/-^ mice^15^.

We found that IGF-1 exerted anti-oxidant and anti-apoptotic effects in FH pigs, which may have contributed to attenuation of atherosclerosis progression, leading to smaller necrotic core size and a more stable plaque phenotype. Indeed, IGF-1 reduced plasma N-tyrosine levels, increased TAC, and concomitantly reduced the number of plaque TUNEL^+^ and pH2A.X^+^ cells. Of note these findings are consistent with the ability of IGF-1 to upregulate expression of the anti-oxidant enzyme glutathione peroxidase in cultured EC^62^ and to downregulate 12/15-lipoxygenase levels in murine plaques^63^, suggesting that these mechanisms may be relevant to the effects of IGF-1 in the FH pig model. Furthermore, elevated SMC apoptosis has been associated with low IGF-1 expression in human advanced plaques^64^ and IGF-1 receptor activation inhibited oxidized lipid-induced apoptosis in SMC through the PI3-kinase/Akt signaling pathway^65^ indicating its involvement in IGF-1 induced suppression of apoptosis.

Single cell transcriptomic analysis of human atherosclerotic plaques has been reported^66^, but critical information linking changes in cell-specific transcriptome with plaque morphology is missing. ST captures all polyadenylated transcripts and provides an unbiased picture of entire transcriptome changes within a spatial context^23^. Thus, ST encodes positional information onto transcripts before sequencing, allowing visualization of expression of virtually any gene of interest within the original tissue section^23^. There are ST generated spatial atlases of disease-free organs and diseased tissues (cancer, Alzheimer’s disease, amyotrophic lateral sclerosis, and rheumatoid arthritis)^23^. However, there are no published reports using ST with atherosclerotic tissue obtained from animal models or humans. We used FH pig coronary plaque specimens to optimize the ST technology and quality controls obtained at the end of experiment correlated well with the manufacturer’s recommendations. We found good agreement between transcriptomic profiles and protein expression data detected by IHC. ST revealed distinct clusters of gene expression within histologically homogeneous parts of the plaque, e.g., the FC which contained almost exclusively cluster 1 and 2 with a clear boundary between these clusters. Cluster 1 had a gene expression pattern, cellular content, and morphological feature (reduced thickness) suggesting elevated sensitivity of this site to plaque rupture. Further studies will be required to determine the clinical relevance of this finding.

ST combined with cell deconvolution algorithm identified genes differentially expressed in response to IGF-1 in spots enriched in SMC, MF and fibromyocytes. ST identified *FOS* and *FOSB* genes as molecules with the highest IGF-1-induced reduction in expression among all ST spots and SMC-high spots. A recent study showed that c-FOS (protein encoded by the *FOS* gene) was involved in oxidized lipid-induced formation of SMC-derived foam cells. SMC-specific c-FOS deficiency prevented formation of foam cells *in vitro* and suppressed atherosclerosis in HFD-fed Apoe^-/-^ mice^67^ demonstrating a fundamental pro-atherogenic role of FOS. *FOS* and *FOSB* encode leucine zipper proteins that dimerize with JUN proteins, thereby forming the transcription factor complex AP-1. *In vitro* and *in vivo* studies have implicated AP-1 as a critical common inflammatory transcription factor mediating progression of atherogenesis^68^ and have suggested that AP-1 inhibition is a promising strategy to treat atherosclerosis^69^. AP-1 was reported to mediate oxidized lipid-induced CXCL14 upregulation in MF-derived foam cells^46^. Here we report that IGF-1-induced FOS/FOSB downregulation correlated with CXCL14 gene expression decrease in MF-high spots. CXCL14 is known to bind and activate another pro-atherogenic cytokine CXCL12^70^. We reported recently that MF-specific IGF-1 overexpression decreased CXCL12 plaque and circulating levels and promoted atherosclerosis in mice^39^. Of note, in the current study we found that IGF-1 markedly decreased circulating CXCL12. Taken together these results suggest that downregulation of the AP-1 complex and suppression of the CXCL14/CXCL12 axis are potential mechanisms contributing to IGF-1-induced anti-atherogenic effects.

Most acute coronary events are related to rupture or erosion of atherosclerotic plaques that are not hemodynamically significant^71^. Thus, plaque stability is a critical determinant of clinical events. Stable plaques are characterized by increased collagen, reduced apoptosis, thicker FC, smaller necrotic cores, and less inflammatory cells^51^. We have shown that coronary plaques in IGF-1-injected FH pigs have reduced necrotic cores, increased FC thickness, decreased levels of MF-like cells and reduced cell apoptosis and these changes are consistent with promotion of a stable plaque phenotype. We also found that IGF-1 decreased the number of MF-high spots, reduced gene expression of CXCL14 chemokine, and downregulated MMP9 gene expression specifically in cluster 1, a potentially more vulnerable component of the FC. Taken together these results suggest that IGF-1 has significant plaque stabilizing properties.

In summary, we report here that IGF-1 administered over 6 months decreased coronary atherosclerosis and promoted features of stable atheroma in FH pigs, without evidence of a tumorigenic effect. IGF-1 stimulated an array of potentially anti-atherogenic mechanisms, including suppression of oxidative stress, systemic inflammatory response, and cell apoptosis. Furthermore IGF-1 decreased EC damage in coronary plaques. ST technology combined with cell deconvolution analysis identified an area of the FC with potentially more propensity for erosion or rupture. Furthermore, ST revealed that IGF-1 induced major changes in the plaque transcriptome. IGF-1 dramatically suppressed gene expression of FOS/FOSB transcription factors and of CXCL14 chemokine and MMP9 in plaques and these data suggest involvement of these molecules in mediating IGF-1 effects. Our results provide mechanistic insights into IGF-1-induced effects on atherosclerosis, are a critical step in taking IGF-1 to human studies, and to our knowledge represent the first report on the use of ST to analyze atherosclerotic tissue from animals or humans.

## ACKNOWLEDGEMENTS

Study was supported by grants from NIH/NHLBI (R01HL070241 and R01HL070241-AS for PD, R01HL142796 for SS). We thanks to Justin Schwarzlose, Jessica Taylor, Vanessa Brown, Jessica Everett, Zhaohui LI (all are University of Missouri-Columbia), Natalya Motyka, Chase Adams, Corey Berta, Ashley Spooner, Kaylynn Fernando, and Nguyen Yen Nhi Ngo (all are Tulane University) for help with pig injections. We appreciate assistance of Ina Nikolli (Harvard University) and Fareed Sindi (Tulane University) with blood and tissue collection. We thank to Andrew Gross (Tulane University) for help with quantification of atherosclerosis and vascular morphology, and Dr. Dae Young Kim (University of Missouri-Columbia) for assistance with necropsy and histopathology. W*e* gratefully acknowledge the help of Kejing Song (Center for Translational Research in Infection and Inflammation, Tulane University) with sequencing of spatial transcriptomics samples.

## Nonstandard Abbreviations and Acronyms

IGF-1: insulin-like growth factor I
CAD: coronary atherosclerotic disease
RCA: right coronary artery
LAD: left anterior descending artery
FH pigs: Rapacz pigs with familial hypercholesterolemia
HFD: high-fat diet
ST: spatial transcriptomics
SMC: smooth muscle cell
MF: macrophages
EC: endothelial cells
FM: fibromyocytes
EEM: external elastic membrane
IEM: internal elastic membrane
TM: tunica media
TA: tunica adventitia
AP: atherosclerotic plaque
NC: necrotic core
FC: fibrous cap
IVUS: intravascular ultrasound
CSA: cross-sectional area
BW: body weight
DEG: differentially expressed genes

## REFERENCES

1. Mozaffarian D, Benjamin EJ, Go AS, Arnett DK, Blaha MJ, Cushman M, Das SR, de Ferranti S, Despres JP, Fullerton HJ, Howard VJ, Huffman MD, Isasi CR, Jimenez MC, Judd SE, Kissela BM, Lichtman JH, Lisabeth LD, Liu S, Mackey RH, Magid DJ, McGuire DK, Mohler ER, 3rd, Moy CS, Muntner P, Mussolino ME, Nasir K, Neumar RW, Nichol G, Palaniappan L, Pandey DK, Reeves MJ, Rodriguez CJ, Rosamond W, Sorlie PD, Stein J, Towfighi A, Turan TN, Virani SS, Woo D, Yeh RW, Turner MB, American Heart Association Statistics C and Stroke Statistics S. Executive Summary: Heart Disease and Stroke Statistics-2016 Update: A Report From the American Heart Association. Circulation. 2016;133:447–54.

2. Tsao CW, Aday AW, Almarzooq ZI, Alonso A, Beaton AZ, Bittencourt MS, Boehme AK, Buxton AE, Carson AP, Commodore-Mensah Y, Elkind MSV, Evenson KR, Eze-Nliam C, Ferguson JF, Generoso G, Ho JE, Kalani R, Khan SS, Kissela BM, Knutson KL, Levine DA, Lewis TT, Liu J, Loop MS, Ma J, Mussolino ME, Navaneethan SD, Perak AM, Poudel R, Rezk-Hanna M, Roth GA, Schroeder EB, Shah SH, Thacker EL, VanWagner LB, Virani SS, Voecks JH, Wang NY, Yaffe K, Martin SS, American Heart Association Council on E, Prevention Statistics C and Stroke Statistics S. Heart Disease and Stroke Statistics-2022 Update: A Report From the American Heart Association. Circulation. 2022:CIR0000000000001052.

3. Arbab-Zadeh A. Coronary Atheroma Burden Is the Main Determinant of Patient Outcome: But How Much Detail Is Needed? Circ Cardiovasc Imaging. 2018;11:e007992.

4. Kaabia Z, Poirier J, Moughaizel M, Aguesse A, Billon-Crossouard S, Fall F, Durand M, Dagher E, Krempf M and Croyal M. Plasma lipidomic analysis reveals strong similarities between lipid fingerprints in human, hamster and mouse compared to other animal species. Sci Rep. 2018;8:15893.

5. Suzuki Y, Yeung AC and Ikeno F. The representative porcine model for human cardiovascular disease. J Biomed Biotechnol. 2011;2011:195483.

6. Dixon JL, Shen S, Vuchetich JP, Wysocka E, Sun GY and Sturek M. Increased atherosclerosis in diabetic dyslipidemic swine: protection by atorvastatin involves decreased VLDL triglycerides but minimal effects on the lipoprotein profile. J Lipid Res. 2002;43:1618–29.

7. Prescott MF, McBride CH, Hasler-Rapacz J, Von Linden J and Rapacz J. Development of complex atherosclerotic lesions in pigs with inherited hyper-LDL cholesterolemia bearing mutant alleles for apolipoprotein B. Am J Pathol. 1991;139:139–47.

8. Schwartz CJ, Valente AJ and Sprague EA. A modern view of atherogenesis. Am J Cardiol. 1993;71:9B–14B.

9. Hasler-Rapacz JO, Nichols TC, Griggs TR, Bellinger DA and Rapacz J. Familial and diet-induced hypercholesterolemia in swine. Lipid, ApoB, and ApoA-I concentrations and distributions in plasma and lipoprotein subfractions. Arterioscler Thromb. 1994;14:923–30.

10. Hamamdzic D and Wilensky RL. Porcine models of accelerated coronary atherosclerosis: role of diabetes mellitus and hypercholesterolemia. J Diabetes Res. 2013;2013:761415.

11. Ross R. Growth factors in the pathogenesis of atherosclerosis. Acta Med Scand Suppl. 1987;715:33–8.

12. Ruotolo G, Bavenholm P, Brismar K, Efendic S, Ericsson CG, de Faire U, Nilsson J and Hamsten A. Serum insulin-like growth factor-I level is independently associated with coronary artery disease progression in young male survivors of myocardial infarction: beneficial effects of bezafibrate treatment. J Am Coll Cardiol. 2000;35:647–54.

13. von der Thusen JH, Borensztajn KS, Moimas S, van Heiningen S, Teeling P, van Berkel TJ and Biessen EA. IGF-1 has plaque-stabilizing effects in atherosclerosis by altering vascular smooth muscle cell phenotype. Am J Pathol. 2011;178:924–34.

14. Higashi Y, Gautam S, Delafontaine P and Sukhanov S. IGF-1 and cardiovascular disease. Growth Horm IGF Res. 2019;45:6–16.

15. Sukhanov S, Higashi Y, Shai SY, Vaughn C, Mohler J, Li Y, Song YH, Titterington J and Delafontaine P. IGF-1 reduces inflammatory responses, suppresses oxidative stress, and decreases atherosclerosis progression in ApoE-deficient mice. Arterioscler Thromb Vasc Biol. 2007;27:2684–90.

16. Schroeter MR, Humboldt T, Schafer K and Konstantinides S. Rosuvastatin reduces atherosclerotic lesions and promotes progenitor cell mobilisation and recruitment in apolipoprotein E knockout mice. Atherosclerosis. 2009;205:63–73.

17. Raber L, Taniwaki M, Zaugg S, Kelbaek H, Roffi M, Holmvang L, Noble S, Pedrazzini G, Moschovitis A, Luscher TF, Matter CM, Serruys PW, Juni P, Garcia-Garcia HM, Windecker S and Investigators IT. Effect of high-intensity statin therapy on atherosclerosis in non-infarct-related coronary arteries (IBIS-4): a serial intravascular ultrasonography study. Eur Heart J. 2015;36:490–500.

18. Juul A, Scheike T, Davidsen M, Gyllenborg J and Jorgensen T. Low serum insulin-like growth factor I is associated with increased risk of ischemic heart disease: a population-based case-control study. Circulation. 2002;106:939–44.

19. Schuler-Luttmann S, Monnig G, Enbergs A, Schulte H, Breithardt G, Assmann G, Kerber S and von Eckardstein A. Insulin-like growth factor-binding protein-3 is associated with the presence and extent of coronary arteriosclerosis. Arterioscler Thromb Vasc Biol. 2000;20:E10–5.

20. Kawachi S, Takeda N, Sasaki A, Kokubo Y, Takami K, Sarui H, Hayashi M, Yamakita N and Yasuda K. Circulating insulin-like growth factor-1 and insulin-like growth factor binding protein-3 are associated with early carotid atherosclerosis. Arterioscler Thromb Vasc Biol. 2005;25:617–21.

21. Lee DL, Wamhoff BR, Katwa LC, Reddy HK, Voelker DJ, Dixon JL and Sturek M. Increased endothelin-induced Ca2+ signaling, tyrosine phosphorylation, and coronary artery disease in diabetic dyslipidemic Swine are prevented by atorvastatin. J Pharmacol Exp Ther. 2003;306:132–40.

22. Shanmugalingam T, Bosco C, Ridley AJ and Van Hemelrijck M. Is there a role for IGF-1 in the development of second primary cancers? Cancer Med. 2016;5:3353–3367.

23. Rao A, Barkley D, Franca GS and Yanai I. Exploring tissue architecture using spatial transcriptomics. Nature. 2021;596:211–220.

24. Butler A, Hoffman P, Smibert P, Papalexi E and Satija R. Integrating single-cell transcriptomic data across different conditions, technologies, and species. Nat Biotechnol. 2018;36:411–420.

25. Lystig TC. Adjusted P values for genome-wide scans. Genetics. 2003;164:1683–7.

26. Fintini D, Brufani C and Cappa M. Profile of mecasermin for the long-term treatment of growth failure in children and adolescents with severe primary IGF-1 deficiency. Ther Clin Risk Manag. 2009;5:553–9.

27. Tavakkol A, Simmen FA and Simmen RC. Porcine insulin-like growth factor-I (pIGF-I): complementary deoxyribonucleic acid cloning and uterine expression of messenger ribonucleic acid encoding evolutionarily conserved IGF-I peptides. Mol Endocrinol. 1988;2:674–81.

28. Stary HC, Chandler AB, Dinsmore RE, Fuster V, Glagov S, Insull W, Jr., Rosenfeld ME, Schwartz CJ, Wagner WD and Wissler RW. A definition of advanced types of atherosclerotic lesions and a histological classification of atherosclerosis. A report from the Committee on Vascular Lesions of the Council on Arteriosclerosis, American Heart Association. Circulation. 1995;92:1355–74.

29. Sukhanov S, Higashi Y, Shai SY, Blackstock C, Galvez S, Vaughn C, Titterington J and Delafontaine P. Differential requirement for nitric oxide in IGF-1-induced anti-apoptotic, anti-oxidant and anti-atherosclerotic effects. FEBS Lett. 2011;585:3065–72.

30. Chakraborty R, Chatterjee P, Dave JM, Ostriker AC, Greif DM, Rzucidlo EM and Martin KA. Targeting smooth muscle cell phenotypic switching in vascular disease. JVS Vasc Sci. 2021;2:79–94.

31. Kovacic JC, Dimmeler S, Harvey RP, Finkel T, Aikawa E, Krenning G and Baker AH. Endothelial to Mesenchymal Transition in Cardiovascular Disease: JACC State-of-the-Art Review. J Am Coll Cardiol. 2019;73:190–209.

32. Blackstock CD, Higashi Y, Sukhanov S, Shai SY, Stefanovic B, Tabony AM, Yoshida T and Delafontaine P. Insulin-like growth factor-1 increases synthesis of collagen type I via induction of the mRNA-binding protein LARP6 expression and binding to the 5’ stem-loop of COL1a1 and COL1a2 mRNA. J Biol Chem. 2014;289:7264–74.

33. Shai SY, Sukhanov S, Higashi Y, Vaughn C, Kelly J and Delafontaine P. Smooth muscle cell-specific insulin-like growth factor-1 overexpression in Apoe-/-mice does not alter atherosclerotic plaque burden but increases features of plaque stability. Arterioscler Thromb Vasc Biol. 2010;30:1916–24.

34. Delafontaine P, Song YH and Li Y. Expression, regulation, and function of IGF-1, IGF-1R, and IGF-1 binding proteins in blood vessels. Arterioscler Thromb Vasc Biol. 2004;24:435–44.

35. Martinet W, Knaapen MW, De Meyer GR, Herman AG and Kockx MM. Oxidative DNA damage and repair in experimental atherosclerosis are reversed by dietary lipid lowering. Circ Res. 2001;88:733–9.

36. Mah LJ, El-Osta A and Karagiannis TC. gammaH2AX: a sensitive molecular marker of DNA damage and repair. Leukemia. 2010;24:679–86.

37. Paffen E and DeMaat MP. C-reactive protein in atherosclerosis: A causal factor? Cardiovasc Res. 2006;71:30–9.

38. Paul A, Ko KW, Li L, Yechoor V, McCrory MA, Szalai AJ and Chan L. C-reactive protein accelerates the progression of atherosclerosis in apolipoprotein E-deficient mice. Circulation. 2004;109:647–55.

39. Snarski P, Sukhanov S, Yoshida T, Higashi Y, Danchuk S, Chandrasekar B, Tian D, Rivera-Lopez V and Delafontaine P. Macrophage-Specific IGF-1 Overexpression Reduces CXCL12 Chemokine Levels and Suppresses Atherosclerotic Burden in Apoe-Deficient Mice. Arterioscler Thromb Vasc Biol. 2022;42:113–126.

40. Kologrivova I, Suslova T, Koshelskaya O, Trubacheva O, Haritonova O and Vinnitskaya I. Frequency of monocyte subsets is linked to the severity of atherosclerosis in patients with ischemic heart disease: A case-control study. Biomedicine (Taipei). 2020;10:36–47.

41. Schroeder A, Mueller O, Stocker S, Salowsky R, Leiber M, Gassmann M, Lightfoot S, Menzel W, Granzow M and Ragg T. The RIN: an RNA integrity number for assigning integrity values to RNA measurements. BMC Mol Biol. 2006;7:3.

42. Dong R and Yuan GC. SpatialDWLS: accurate deconvolution of spatial transcriptomic data. Genome Biol. 2021;22:145.

43. Wirka RC, Wagh D, Paik DT, Pjanic M, Nguyen T, Miller CL, Kundu R, Nagao M, Coller J, Koyano TK, Fong R, Woo YJ, Liu B, Montgomery SB, Wu JC, Zhu K, Chang R, Alamprese M, Tallquist MD, Kim JB and Quertermous T. Atheroprotective roles of smooth muscle cell phenotypic modulation and the TCF21 disease gene as revealed by single-cell analysis. Nat Med. 2019;25:1280–1289.

44. Lafontaine DL and Tollervey D. The function and synthesis of ribosomes. Nat Rev Mol Cell Biol. 2001;2:514–20.

45. Snarski P, Sukhanov S, Yoshida T, Higashi Y, Danchuk S, Chandrasekar B, Tian D, Rivera-Lopez V and Delafontaine P. Macrophage-Specific IGF-1 Overexpression Reduces CXCL12 Chemokine Levels and Suppresses Atherosclerotic Burden in Apoe-Deficient Mice. Arterioscler Thromb Vasc Biol. 2021:ATVBAHA121316090.

46. Tong W, Duan Y, Yang R, Wang Y, Peng C, Huo Z and Wang G. Foam Cell-Derived CXCL14 Muti-Functionally Promotes Atherogenesis and Is a Potent Therapeutic Target in Atherosclerosis. J Cardiovasc Transl Res. 2020;13:215–224.

47. Perbal B. CCN proteins: A centralized communication network. J Cell Commun Signal. 2013;7:169–77.

48. To WS and Midwood KS. Plasma and cellular fibronectin: distinct and independent functions during tissue repair. Fibrogenesis Tissue Repair. 2011;4:21.

49. Zhao CF and Herrington DM. The function of cathepsins B, D, and X in atherosclerosis. Am J Cardiovasc Dis. 2016;6:163–170.

50. Eckert J, Schmidt M, Magedanz A, Voigtlander T and Schmermund A. Coronary CT angiography in managing atherosclerosis. Int J Mol Sci. 2015;16:3740–56.

51. Kolodgie FD, Gold HK, Burke AP, Fowler DR, Kruth HS, Weber DK, Farb A, Guerrero LJ, Hayase M, Kutys R, Narula J, Finn AV and Virmani R. Intraplaque hemorrhage and progression of coronary atheroma. N Engl J Med. 2003;349:2316–25.

52. Man JJ, Beckman JA and Jaffe IZ. Sex as a Biological Variable in Atherosclerosis. Circ Res. 2020;126:1297–1319.

53. Link RP, Pedersoli WM and Safanie AH. Effect of exercise on development of atherosclerosis in swine. Atherosclerosis. 1972;15:107–22.

54. Kobari Y, Koto M and Tanigawa M. Regression of diet-induced atherosclerosis in Gottingen miniature swine. Lab Anim. 1991;25:110–6.

55. Oppi S, Luscher TF and Stein S. Mouse Models for Atherosclerosis Research-Which Is My Line? Front Cardiovasc Med. 2019;6:46.

56. Fan J, Kitajima S, Watanabe T, Xu J, Zhang J, Liu E and Chen YE. Rabbit models for the study of human atherosclerosis: from pathophysiological mechanisms to translational medicine. Pharmacol Ther. 2015;146:104–19.

57. Hajar R. Risk Factors for Coronary Artery Disease: Historical Perspectives. Heart Views. 2017;18:109–114.

58. Davignon J and Ganz P. Role of endothelial dysfunction in atherosclerosis. Circulation. 2004;109:III27–32.

59. Sluiter TJ, van Buul JD, Huveneers S, Quax PHA and de Vries MR. Endothelial Barrier Function and Leukocyte Transmigration in Atherosclerosis. Biomedicines. 2021;9.

60. Higashi Y, Sukhanov S, Shai SY, Danchuk S, Snarski P, Li Z, Hou X, Hamblin MH, Woods TC, Wang M, Wang D, Yu H, Korthuis RJ, Yoshida T and Delafontaine P. Endothelial deficiency of insulin-like growth factor-1 receptor reduces endothelial barrier function and promotes atherosclerosis in Apoe-deficient mice. Am J Physiol Heart Circ Physiol. 2020;319:H730–H743.

61. Bach LA. Endothelial cells and the IGF system. J Mol Endocrinol. 2015;54:R1–13.

62. Higashi Y, Pandey A, Goodwin B and Delafontaine P. Insulin-like growth factor-1 regulates glutathione peroxidase expression and activity in vascular endothelial cells: Implications for atheroprotective actions of insulin-like growth factor-1. Biochim Biophys Acta. 2013;1832:391–9.

63. Sukhanov S, Snarski P, Vaughn C, Lobelle-Rich P, Kim C, Higashi Y, Shai SY and Delafontaine P. Insulin-like growth factor I reduces lipid oxidation and foam cell formation via downregulation of 12/15-lipoxygenase. Atherosclerosis. 2015;238:313–20.

64. Okura Y, Brink M, Zahid AA, Anwar A and Delafontaine P. Decreased expression of insulin-like growth factor-1 and apoptosis of vascular smooth muscle cells in human atherosclerotic plaque. J Mol Cell Cardiol. 2001;33:1777–89.

65. Li Y, Higashi Y, Itabe H, Song YH, D. J and Delafontaine P. Insulin-like growth factor-1 receptor activation inhibits oxidized LDL-induced cytochrome C release and apoptosis via the phosphatidylinositol 3 kinase/Akt signaling pathway. Arterioscler Thromb Vasc Biol. 2003;23:2178–84.

66. Depuydt MAC, Prange KHM, Slenders L, Ord T, Elbersen D, Boltjes A, de Jager SCA, Asselbergs FW, de Borst GJ, Aavik E, Lonnberg T, Lutgens E, Glass CK, den Ruijter HM, Kaikkonen MU, Bot I, Slutter B, van der Laan SW, Yla-Herttuala S, Mokry M, Kuiper J, de Winther MPJ and Pasterkamp G. Microanatomy of the Human Atherosclerotic Plaque by Single-Cell Transcriptomics. Circ Res. 2020;127:1437–1455.

67. Miao G, Zhao X, Chan SL, Zhang L, Li Y, Zhang Y, Zhang L and Wang B. Vascular smooth muscle cell c-Fos is critical for foam cell formation and atherosclerosis. Metabolism. 2022;132:155213.

68. Ye N, Ding Y, Wild C, Shen Q and Zhou J. Small molecule inhibitors targeting activator protein 1 (AP-1). J Med Chem. 2014;57:6930–48.

69. Cheng SM, Yang SP, Ho LJ, Tsao TP, Chang DM and Lai JH. Irbesartan inhibits human T-lymphocyte activation through downregulation of activator protein-1. Br J Pharmacol. 2004;142:933–42.

70. Kouzeli A, Collins PJ, Metzemaekers M, Meyrath M, Szpakowska M, Artinger M, Struyf S, Proost P, Chevigne A, Legler DF, Eberl M and Moser B. CXCL14 Preferentially Synergizes With Homeostatic Chemokine Receptor Systems. Front Immunol. 2020;11:561404.

71. Ambrose JA, Tannenbaum MA, Alexopoulos D, Hjemdahl-Monsen CE, Leavy J, Weiss M, Borrico S, Gorlin R and Fuster V. Angiographic progression of coronary artery disease and the development of myocardial infarction. J Am Coll Cardiol. 1988;12:56–62.

