## Supplemental Methods, Supplemental Figure Legends for "INSULIN-LIKE GROWTH FACTOR I REDUCES CORONARY ATHEROSCLEROSIS IN PIGS WITH FAMILIAL HYPERCHOLESTEROLEMIA"

**SUPLEMENTAL MATERIALS**

**EXTENDED METHODS**

***Intravascular ultrasound (IVUS).*** We assigned 2 pigs per IVUS procedure/day: one pig from the IGF-1 group and one from the saline group. Pigs were pre-treated with a daily dose of aspirin (325 mg) for 3 days before IVUS. All animals were fasted for 12 hours prior to IVUS procedure. Telazol (2.25-6 mg/kg IM) and xylazine (1-2.25 mg/kg IM) were administered for anesthesia. An intravenous catheter was placed in a marginal ear vein for the administration of fluids and drugs. The auricular artery was canulated to allow for arterial blood pressure measurements and baseline blood pressure. Diazepam (0.5-1.0 mg/kg IV) was given to aid with intubation. An appropriately sized endotracheal tube was inserted for mechanical ventilation. Pigs were intubated and ventilated with oxygen mixed with nitrogen (compressed medical air). Ventilation parameters were set to maintain inspiratory pressures between 15-30 mmHg, respiration rate of 10-20 BPM and spO_2_ 92-100%. Arterial blood gases were taken and if pO_2_, CO_2_ and pH were out of the reference range, ventilation parameters were adjusted to correct. General anesthesia was maintained with isoflurane (1.5 to 3%) in O_2_. Blood oxygenation was monitored using continuous pulse oximetry. Electrocardiogram was monitored throughout the entire procedure to detect myocardial infarction and possible arrhythmias requiring medical intervention. The pig was placed on a heating blanket to maintain rectal temperature at 36.5 to 39.0°C (assessed via rectal probe). Artificial tears ophthalmic ointment was applied bilaterally to the corneal surfaces to keep the corneas moist during the procedure. After intubation and clipping the vascular access area, a sterile prep was performed using 3 cycles of surgical scrub betadine or chlorohexidine. The right or left femoral artery and vein were cannulated with a percutaneous sheath for catheter advancement into the coronary arteries. To prevent thrombosis throughout the duration of the procedure, a loading dose of heparin was administered as needed to achieve an activated clotting time >300 seconds. Contrast-enhanced coronary cine angiographic images were acquired and stored using the Innova Optima CL323i X-ray system (GE Healthcare, Pewaukee, WI). Intravascular ultrasound was performed using the IVUS imaging system (Volcano Corporation, San Diego, CA) and a 20 MHz 3.5 F Visions PV 0.035 Digital IVUS Catheter (Volcano Corporation, San Diego, CA) advanced to the left anterior descending artery (LAD) and the right coronary artery (RCA) through a standard 5 F guide catheter. Operative management included fluid maintenance (0.9% saline) and the administration of long-acting cefazolin sodium (15 mg/kg, IM) as antibiotic prophylaxis. After the procedure, the catheters were removed, and the artery closed (Angio-Seal Vascular Closure Device, Terumo Medical Corp, New Jersey, NJ). Analgesia was provided with Buprenex-SR (0.12-0.27 mg/kg SC). IVUS was performed at baseline (T0) and at 3 and 6 months (T3, T6, Fig1A)). Following the final invasive imaging procedure, pigs were euthanized while under deep isoflurane anesthesia using potassium chloride (KCl, 20 mEq IV) to arrest the heart in diastole.

To inject solutions, we used a Stratis needle-free injection system (PharmaJet, Golden, CO). All pigs received a high-fat diet (HFD) starting the day after the T0 IVUS. The HFD contains 800 g/day (by weight 13% protein, 39.6% carbohydrate, 47.4% fat and 2% cholesterol)^1^. Pigs have free access to water. Body weight was measured weekly.

***Blood biochemistry.*** Fasting blood samples were collected from the jugular vein at baseline and monthly. Blood was collected in 0.1 mol/L citrate containing EDTA. Fresh whole blood was submitted to the Antech Diagnostics (New Orleans, LA) for complete blood count (CBC) with differential, and biochemistry measurements (Superchem w/CBC cat# SA020). Plasma IGF-1 levels were quantified by human IGF-I Quantikine ELISA kit (cat# DG100B) and C-reactive protein levels by porcine C-reactive protein/CRP DuoSet ELISA (cat# DY2648, both from R&D Systems). Quantification of plasma N-tyrosine and total antioxidant capacity (TAC) assay were performed with OxiSelect™ Nitrotyrosine ELISA Kit (cat# STA-305) and TAC assay kit (cat# STA-360) (both from Cell Biolabs Inc, San Diego, CA). TAC assay measures total antioxidant capacity based on reduction of copper (II) to copper (I).

***IVUS analysis.*** Coronary arteries contain the tunica intima layer (a complex of endothelium, atheroma, and internal elastic membrane (IEM)), tunica media layer (TM) (TM consists of smooth muscle and external elastic membrane (EEM)) and tunica externa layer which comprises the tunica adventitia (TA) and peri-adventitial tissues^2^. Since the leading edge of the media (corresponding to the IEM) is not well delineated, IVUS measurements cannot determine true histological atheroma area. Consistent with the 2021 clinical expert consensus document on IVUS measurements the EEM and lumen areas are used to calculate a surrogate for true atherosclerotic plaque area and the term “plaque plus media area” is recommended^2^. For the current study 20 mm of IVUS pullback segment distal to the ostia of the right coronary artery (RCA) and the left anterior descending artery (LAD) were selected. The area circumscribed by the outer border of the echolucent TM and the luminal border was manually traced on each 1 mm IVUS frame within selected fragment. SS and ZR independently performed the manual outline of both areas, and ZR was blinded to the individual animal assignment. The inter-rater variability was 10.5%, 7.2% and 10.8% for evaluation of lumen volume, vessel volume and relative atheroma volume, respectively.

***Atherosclerotic burden assessment by histology.*** Following completion of the experimental protocol, hearts were excised in accordance with the recommendation of the American Veterinary Medical Association Guide on Euthanasia. The entire right coronary (RCA) and the left anterior descending (LAD) arteries were collected, and the proximal 30 mm fragment of RCA and LAD was immersed into 10% formalin, and further cut onto 5 mm fragments. One Trichrome-stained section per each block was used to quantify atherosclerotic burden. Sections were imaged with Olympus IX71 inverted microscope equipped with DP80 camera. Analysis of vessel morphometry was performed with CellSens Dimension 1.18 software (Olympus) by two independent researchers, one of them blinded. The mean±SEM discrepancy between evaluators was 11.2±1.7%. EEM, IEM and luminal border were manually outlined, and corresponding cross-sectional areas (CSA) were measured.

The following indices were assessed:

1) relative tunica media CSA = (EEM-IEM)/EEM x 100%.
2) relative atherosclerotic plaque CSA = (IEM-Luminal area)/EEM x 100%.
3) necrotic core (NC) area and fibrous cap (FC) area were manually outlined using CellSens Dimension software. The NC was defined as acellular (hematoxylin-negative) plaque area.
4) FC was defined as largely uninterrupted strip of brown-colored (smooth muscle, connective tissues) material on the top of necrotic or lipid core with a higher density of nuclei than the plaque core. The thickness of FC was calculated as the mean length of 5 arbitrary lines distributed across the cap area.

***Plaque composition analysis.*** The cellular content of coronary plaques was assessed by immunohistochemistry (IHC). To perform IHC coronary sections were deparaffinized, dehydrated and processed with heat-mediated antigen retrieval using citrate buffer (pH 6.0) followed by blocking step (Protein block, Abcam, cat# ab64226). Sections were incubated overnight at +4^o^C with mixture of primary antibodies, or with mixture of normal IgG (negative control). The 1^st^ section was incubated with mouse α-SMA antibody (Millipore, cat# CBL171, clone ASM1) plus rabbit pH2A.X histone antibody (Cell Signaling, cat# 9718, clone 20E3). The 2^nd^ serial section was incubated with mouse SRA antibody (Trans Genia Inc., cat# KT022, clone SRA-E5) plus rabbit CD31 antibody (Abcam, cat# 134168, clone EP3095) and the 3^rd^ serial section with mouse IgG plus rabbit IgG (both are from Santa Cruz Biotechnology, cat# sc-2025 and sc-2027, respectively). Rabbit antibody signal was visualized with goat anti-rabbit-biotin IgG (Abcam, cat# BA-1000) followed by incubation with streptavidin-AlexaFluor594 conjugate (Life Technologies, cat# S32356) plus DAPI. Mouse antibody signal was amplified with Alexa Fluor 488 Tyramide Super Boost kit (Invitrogen, cat# B40912). Sections were mounted with ProLong Gold antifade media (Thermo Fisher, cat# P36970) for imaging. To quantify immunopositivity for each antibody, sections were scanned with Cytation 5 multi-mode imager (Bio-Tek, Winooski, MI) using standard Texas Red, GFP and DAPI filter cubes to generate greyscale images for each channel. Next, vessel morphological regions-of-interest (ROI) (i.e., TM, plaque, etc.) were manually outlined and area positive for cellular marker was quantified within ROI. The ratio of marker-positive area per ROI area (x 100%) was calculated and shown in Figures.

SMC and MF share cell markers in the atherosclerotic plaque^3^ and plaque EC undergo a change in phenotype toward a mesenchymal cell type^4^. Such phenotype switching complicates marker-based cell identification. To validate the IHC protocol, serial RCA sections were stained with a set of cell marker antibodies and immunopositivity pattern was compared. We found that each of 4 SMC marker antibodies stained virtually identical cell population in the plaque and a similar conclusion was made for 3 tested MF and 3 EC marker antibodies (Suppl.Fig.2). These data show that IHC with antibody for a single cell marker identifies plaque cells expressing multiple markers increasing confidence to identify specific plaque cells. We confirmed that cells immunopositive for SRA, a MF marker were immunonegative for α-SMA, a SMC marker and *vice versa* (Suppl.Fig.2A) showing that these antibodies have no cross-reactivity. IHC for proliferating cell nuclear antigen (PCNA, marker of cell proliferation^5^ was performed with mouse anti-PCNA antibody (cat# MAB424R, clone PC10, Millipore-Sigma) using anti-mouse-biotin IgG (Thermo Fisher, cat# 31800) followed by incubation with streptavidin-AlexaFluor594 conjugate plus DAPI.

***Cell apoptosis.*** To quantify cell apoptosis, we used *In Situ* Cell Death Detection Kit, TMR red (cat# 12156792910, Millipore-Sigma**)** as per manufacturer’s instructions. Briefly, sections were deparaffinized, dehydrated and permeabilized with Digest-All 2 (Trypsin) kit (Invitrogen, cat# OO3008). Next, sections were incubated with TUNEL mixture diluted 1:2 with TUNEL dilution buffer (cat # 11966006001, Roche) for 45 min at 37^o^C, washed with PBS, co-stained with DAPI and mounted with ProLong Gold antifade media. Treatment with DNase I (Ambion, cat# AM2222, 10U/sections) for 20 min at 37^o^C was used to generate positive control. Each slide with TUNEL-stained section contained a serial section stained with dUTP-omitted TUNEL mixture serving as a negative control. Total cell apoptosis was defined as TUNEL-positive cell number per 1000 DAPI-positive nuclei.

***Monocyte subsets.*** To count circulating CD163^lo^/CD14^hi^ monocytes, whole blood was mixed with a cocktail of monoclonal antibodies against CD163-PE (1: 80 dilution, Bio-Rad, cat# MCA2311PE), and CD14-Alexa Fluor 488 (1: 20 dilution, Bio-Rad, cat# MCA1568A488), porcine CD172a (1: 100 dilution, Southern Biotech, cat# 4525-08), or appropriate isotype controls and subsequently with streptavidin-APC/Cy7 (1: 1000 dilution, Southern Biotech, cat# 7105-19). Red blood cells were lysed by BD Pharm Lyse™ lysing solution (BD Biosciences, cat# 555899) and washed 3 times using PBS before flow-cytometric analysis. Data acquisition and analysis were performed using Beckman Coulter Epics Gallios analyzer running Gallios software v1.1; CD172a-positive leukocytes were further size-gated to identify monocytes. The monocyte-gate was differentiated into two subsets based on CD163 and CD14 expression levels and counted as CD163^lo^/CD14^hi^ monocytes and CD163^hi^/CD14^lo^ monocytes^6^.

***Spatial transcriptomics (ST).*** ST utilizes spotted arrays of specialized mRNA-capturing probes containing a spatial barcode unique to that spot. When a cryosection is attached to the slide, the capture probes bind mRNA from the adjacent point in the tissue. After mRNA extraction, cDNA library is generated and sequenced. ST was conducted using the Visium Spatial Gene Expression System (10× Genomics, Pleasanton, CA). The RCA fragments were snap-frozen, embedded with OCT compound and cut at –26°C at a thickness of 10 µm. Optimization and gene expression assays were carried out according to the manufacturer’s protocol. Briefly, slides were fixed in –20°C methanol, dried with isopropanol, and stained with H&E. A tile scan image of the entire section was generated using a Cytation 5 imager. For tissue optimization, enzymatic permeabilization was conducted for 0–40 min, followed by first-strand cDNA synthesis with fluorescent nucleotides. The slide was reimaged using standard RFP filter cube. An optimal permeabilization time of 12 min was determined by visual inspection to maximize mRNA recovery while at the same time minimizing diffusion. Library preparation, clean-up, and indexing were conducted using company manual-guided procedures. Samples were subjected to pair-ended sequencing using an Illumina NextSeq 2000 generating ~800 M reads. Pig reference genome was created from *Sus scrofa* genomic sequence (Sscrofa 11.1) and Ensembl annotation, and reads were aligned and counted by Space Ranger (10x Genomics). All the downstream analyses were performed using R toolkit Seurat^7^ and Ingenuity pathway analysis (IPA) software (Qiagen). SMC-, MF- and fibromyocytes (FM) distribution and calculation of cell type ratio were predicted by mixed cell deconvolution analysis driven by Seurat using human atherosclerotic RCA single cell RNA seq dataset (accession number GSE131778)^8^. The accession number for the deposition of ST data is pending.

**AUTHORS CONTRIBUTION**

**SS designed the study, performed histological assays, IVUS analysis, ran spatial transcriptomic and wrote the manuscript; YH designed the study, performed blood biochemistry and monocyte subsets assessment and contributed to writing the manuscript; SD and MA carried out ELISAs, TY and JK handled spatial transcriptomic data and ran bioinformatics, TG, AS, TS, JJ, DG, JI, DT, and DB performed IVUS surgery; JS carried out necropsy, ZR assisted with IVUS analysis, DB, DL and PD designed the study and contributed to writing the manuscript.**

**SUPPLEMENTAL FIGURE LEGENDS**

**Suppl.Fig.1. FH pigs body weight, blood pressure and heart rate.** IGF-1 or saline (control) was injected into FH pigs (males, 5/group, females, 9/group) and pigs were fed with high-fat diet (HFD) for 6 months. Body weight was measured weekly. Blood pressure and heart rate were assessed in sedated pigs during IVUS procedure: at basal level (T0), after 3 months of injections (T3) and after 6 months (T6, at sacrificing).

**Suppl.Fig.2. Validation of using macrophages (MF, A), smooth muscle cell (SMC, B) and endothelial cell (EC, C) marker antibody for immunohistochemistry.** RCA serial cross-sections were obtained from FH females. A, Sections were immunostained with antibody for macrophage scavenger receptor A (SRA), monocyte chemoattractant protein 1 (MCP-1), lysosome-associated membrane protein 2 (LAMP2) (all are macrophage markers) and co-stained with α-smooth muscle actin (α-SMA) antibody (SMC marker) conjugated to Alexa Fluor 488 (signal shown in green) and DAPI. Macrophage marker antibody were visualized with biotin/streptavidin amplification system using Alexa Fluor 594 as a fluorescent reporter (shown in red). Note that each MF marker antibody stained virtually identical cell population in the plaque and cells immunopositive for SRA are immunonegative for α-SMA, and *vice versa* (upper left panel). Lu, lumen, TM, tunica media, TA, tunica adventitia, FC. Fibrous cap. B, Sections were stained with α-SMA, calponin, smooth muscle protein 22α (SM22α), myosin heavy chain 11 antibody (all are SMC markers). Each of SMC antibody detected identical cell population in the plaque and vascular media. C, Sections were stained with endothelial nitric oxide synthase (eNOS), CD31, VE-cadherin antibody (all are EC markers). Each of EC antibody detected identical cells on the luminal border.

**Suppl.Fig.3. IGF-1 effect on plaque SMC and collagen.** IGF-1 or saline (control) was injected into FH pigs and pigs were fed with high-fat diet (HFD). RCA and LAD cross-sections were immunostained with α-smooth muscle actin (α-SMA) antibody (SMC marker) (A) or stained with Gomori’s Trichrome stain to visualize collagen (B, C). Sections were imaged and tunica media (TM) and atherosclerotic plaque cross-sectional area were manually outlined. M, males, F, females.

**Suppl.Fig.4. Spatial transcriptomics (ST) clustering.** RCA cryosections from IGF-1- and saline-injected FH females (N=2/group) were processed with ST protocol. All ST spots in IGF-1 and saline specimens were grouped into 9 clusters (0-8) based on their transcriptome. In parallel, plaque’s fibrous cap, lipid core, tunica media and tunica adventitia were outlined using H&E image and transcriptome clusters and histological annotations were compared side-by-side. A, cell type ratio (%) was calculated for each ST spot to identify spots enriched by SMC, MF, fibromyocytes (FM), B cells, T cells, epithelial cells, and fibroblasts. B, cell type ratio for each transcriptome cluster shown in Figure.

**REFERENCES**

1.	Masseau I and Bowles DK. Carotid Endothelial VCAM-1 Is an Early Marker of Carotid Atherosclerosis and Predicts Coronary Artery Disease in Swine. *J Biomed Sci Eng*. 2015;8:789-796.

2.	Saito Y, Kobayashi Y, Fujii K, Sonoda S, Tsujita K, Hibi K, Morino Y, Okura H, Ikari Y and Honye J. Clinical expert consensus document on intravascular ultrasound from the Japanese Association of Cardiovascular Intervention and Therapeutics (2021). *Cardiovasc Interv Ther*. 2022;37:40-51.

3.	Chakraborty R, Chatterjee P, Dave JM, Ostriker AC, Greif DM, Rzucidlo EM and Martin KA. Targeting smooth muscle cell phenotypic switching in vascular disease. *JVS Vasc Sci*. 2021;2:79-94.

4.	Kovacic JC, Dimmeler S, Harvey RP, Finkel T, Aikawa E, Krenning G and Baker AH. Endothelial to Mesenchymal Transition in Cardiovascular Disease: JACC State-of-the-Art Review. *J Am Coll Cardiol*. 2019;73:190-209.

5.	Strzalka W and Ziemienowicz A. Proliferating cell nuclear antigen (PCNA): a key factor in DNA replication and cell cycle regulation. *Ann Bot*. 2011;107:1127-40.

6.	Gordon S and Taylor PR. Monocyte and macrophage heterogeneity. *Nat Rev Immunol*. 2005;5:953-64.

7.	Butler A, Hoffman P, Smibert P, Papalexi E and Satija R. Integrating single-cell transcriptomic data across different conditions, technologies, and species. *Nat Biotechnol*. 2018;36:411-420.

8.	Wirka RC, Wagh D, Paik DT, Pjanic M, Nguyen T, Miller CL, Kundu R, Nagao M, Coller J, Koyano TK, Fong R, Woo YJ, Liu B, Montgomery SB, Wu JC, Zhu K, Chang R, Alamprese M, Tallquist MD, Kim JB and Quertermous T. Atheroprotective roles of smooth muscle cell phenotypic modulation and the TCF21 disease gene as revealed by single-cell analysis. *Nat Med*. 2019;25:1280-1289.
