## Supplemental Figures for "INSULIN-LIKE GROWTH FACTOR I REDUCES CORONARY ATHEROSCLEROSIS IN PIGS WITH FAMILIAL HYPERCHOLESTEROLEMIA"

### Supplemental Figure 1

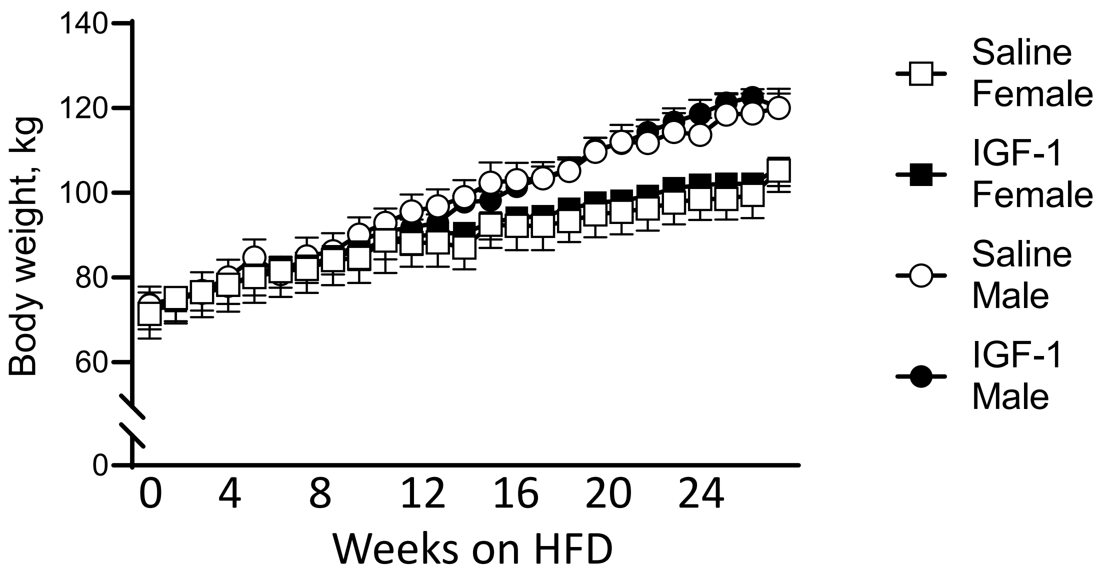


Supplemental Table 1. FH pigs blood pressure (BP) and heart rate (HR)

| Time-point/drug | Sex | BP, mmHg, systolic | | BP, mmHg, diastolic | | HR, bpm | |
|---|---|---|---|---|---|---|---|
| | | **Mean** | **SEM** | **Mean** | **SEM** | **Mean** | **SEM** |
| **T0** | M | 109 | 5.3 | 83.6 | 6.7 | 93.6 | 7.8 |
| | F | 114.5 | 6.6 | 76.3 | 4.5 | 102.7 | 5.8 |
| **T3/Saline** | M | 118.0 | 6.6 | 91.0 | 3.8 | 104.3 | 7.3 |
| | F | 130.7 | 14.5 | 87.2 | 8.0 | 94.0 | 12.9 |
| **T3/IGF-1** | M | 129.3 | 4.5 | 85.5 | 4.5 | 102.2 | 4.4 |
| | F | 121.8 | 7.6 | 81.0 | 4.0 | 104.7 | 9.8 |
| **T6/Saline** | M | 127.0 | 9.1 | 84.0 | 7.5 | 100.5 | 8.6 |
| | F | 133.7 | 3.7 | 91.8 | 8.6 | 89.0 | 3.9 |
| **T6/IGF-1** | M | 120.2 | 7.8 | 72.0 | 7.1 | 99.3 | 6.1 |
| | F | 116.5 | 7.3 | 68.8 | 4.8 | 100.2 | 6.7 |

### Supplemental Table 2. FH pigs blood biochemistry

| Male | Units | Group | 0 |  | 1-mo |  | 2-mo |  | 3-mo |  | 4-mo |  | 5-mo |  | 6-mo |  |
|---|---|---|---|---|---|---|---|---|---|---|---|---|---|---|---|---|
|  |  |  | Avg | SD | Avg | SD | Avg | SD | Avg | SD | Avg | SD | Avg | SD | Avg | SD |
| Total Protein | g/dL | Saline | 6.5 | 0.4 | 6.7 | 0.3 | 6.3 | 0.4 | 5.8 | 0.3 | 6.2 | 0.3 | 6.4 | 0.4 | 6.5 | 0.4 |
|  |  | IGF-1 | 6.4 | 0.3 | 6.4 | 0.1 | 6.0 | 0.3 | 5.9 | 0.3 | 6.1 | 0.4 | 6.3 | 0.3 | 6.2 | 0.4 |
| Albumin | g/dL | Saline | 4.1 | 0.1 | 4.2 | 0.1 | 4.1 | 0.2 | 3.9 | 0.2 | 4.1 | 0.3 | 4.0 | 0.5 | 4.0 | 0.4 |
|  |  | IGF-1 | 4.0 | 0.2 | 3.8 | 0.1 | 3.5 | 0.1 | 3.5 | 0.1 | 3.7 | 0.2 | 3.9 | 0.2 | 3.7 | 0.2 |
| Globulin | g/dL | Saline | 2.4 | 0.4 | 2.5 | 0.3 | 2.2 | 0.5 | 1.9 | 0.4 | 2.1 | 0.3 | 2.3 | 0.6 | 2.5 | 0.7 |
|  |  | IGF-1 | 2.4 | 0.3 | 2.6 | 0.2 | 2.4 | 0.3 | 2.4 | 0.3 | 2.4 | 0.3 | 2.5 | 0.3 | 2.5 | 0.3 |
| A/G Ratio |  | Saline | 1.8 | 0.4 | 1.7 | 0.2 | 2.0 | 0.4 | 2.1 | 0.5 | 2.0 | 0.4 | 1.9 | 0.7 | 1.7 | 0.6 |
|  |  | IGF-1 | 1.7 | 0.3 | 1.5 | 0.2 | 1.5 | 0.2 | 1.5 | 0.2 | 1.6 | 0.3 | 1.6 | 0.3 | 1.5 | 0.3 |
| AST (SGOT) | IU/L | Saline | 24 | 2.9 | 39 | 17.0 | 50 | 64.6 | 22 | 3.3 | 24 | 3.8 | 26 | 9.4 | 20 | 2.5 |
|  |  | IGF-1 | 30 | 12.4 | 25 | 4.3 | 24 | 6.1 | 23 | 6.6 | 22 | 6.0 | 21 | 3.1 | 28 | 18.4 |
| ALT (SGPT) | IU/L | Saline | 28 | 0.8 | 29 | 3.0 | 28 | 5.5 | 23 | 2.4 | 22 | 2.4 | 20 | 3.3 | 21 | 2.3 |
|  |  | IGF-1 | 33 | 6.6 | 32 | 2.8 | 27 | 2.9 | 24 | 2.0 | 23 | 1.6 | 21 | 2.3 | 27 | 9.8 |
| Alk Phosphatase | IU/L | Saline | 78 | 30 | 96 | 39 | 95 | 44 | 81 | 38 | 93 | 38 | 75 | 28 | 83 | 41 |
|  |  | IGF-1 | 80 | 22 | 107 | 23 | 90 | 17 | 79 | 21 | 80 | 22 | 84 | 23 | 82 | 37 |
| GGT | IU/L | Saline | 28 | 9 | 32 | 10 | 28 | 9 | 28 | 9 | 32 | 10 | 34 | 8 | 31 | 8 |
|  |  | IGF-1 | 34 | 9 | 37 | 8 | 34 | 7 | 32 | 7 | 33 | 5 | 37 | 7 | 38 | 5 |
| Total Bilirubin | mg/dL | Saline | 0.1 | 0.0 | 0.2 | 0.0 | 0.1 | 0.0 | 0.2 | 0.1 | 0.1 | 0.0 | 0.1 | 0.0 | 0.1 | 0.0 |
|  |  | IGF-1 | 0.1 | 0.0 | 0.1 | 0.1 | 0.1 | 0.0 | 0.2 | 0.0 | 0.1 | 0.1 | 0.1 | 0.0 | 0.1 | 0.0 |
| BUN | mg/dL | Saline | 10 | 1.8 | 9 | 1.6 | 10 | 0.8 | 9 | 2.2 | 9 | 1.5 | 12 | 2.6 | 13 | 1.8 |
|  |  | IGF-1 | 9 | 1.3 | 8 | 1.8 | 8 | 2.0 | 8 | 1.7 | 7 | 1.5 | 9 | 1.9 | 12 | 1.8 |
| Creatinine | mg/dL | Saline | 1.5 | 0.1 | 1.5 | 0.1 | 1.6 | 0.2 | 1.6 | 0.1 | 1.4 | 0.2 | 1.5 | 0.1 | 1.5 | 0.2 |
|  |  | IGF-1 | 1.4 | 0.1 | 1.4 | 0.2 | 1.5 | 0.1 | 1.4 | 0.2 | 1.3 | 0.1 | 1.4 | 0.1 | 1.4 | 0.2 |
| BUN/CREAT RATIO |  | Saline | 7 | 1.3 | 6 | 0.9 | 6 | 0.0 | 6 | 0.9 | 7 | 1.3 | 8 | 1.6 | 8 | 1.6 |
|  |  | IGF-1 | 7 | 1.5 | 6 | 1.1 | 5 | 1.7 | 6 | 1.8 | 6 | 1.3 | 6 | 1.2 | 9 | 0.5 |
| Phosphorus | mg/dL | Saline | 6.9 | 0.7 | 6.1 | 0.5 | 6.5 | 0.5 | 5.6 | 0.3 | 5.5 | 0.1 | 5.7 | 0.3 | 6.0 | 0.5 |
|  |  | IGF-1 | 7.4 | 0.9 | 6.5 | 0.8 | 5.7 | 0.8 | 6.2 | 0.8 | 5.8 | 0.7 | 6.3 | 0.4 | 6.3 | 0.6 |
| Glucose | mg/dL | Saline | 156 | 80 | 79 | 12 | 87 | 9 | 180 | 41 | 92 | 7 | 100 | 16 | 132 | 39 |
|  |  | IGF-1 | 90 | 16 | 83 | 7 | 117 | 23 | 108 | 44 | 90 | 9 | 116 | 45 | 133 | 11 |
| CALCIUM | mg/dL | Saline | 10.7 | 0.5 | 10.8 | 0.2 | 10.7 | 0.2 | 10.6 | 0.2 | 10.6 | 0.3 | 10.5 | 0.5 | 10.7 | 0.2 |
|  |  | IGF-1 | 10.6 | 0.5 | 10.5 | 0.2 | 10.2 | 0.1 | 10.3 | 0.5 | 10.3 | 0.1 | 10.3 | 0.3 | 10.2 | 0.3 |
| Magnesium | mEq/L | Saline | 2.0 | 0.3 | 1.9 | 0.2 | 2.0 | 0.2 | 1.9 | 0.1 | 2.0 | 0.2 | 1.9 | 0.1 | 2.0 | 0.2 |
|  |  | IGF-1 | 1.9 | 0.2 | 1.9 | 0.2 | 1.8 | 0.1 | 1.8 | 0.2 | 1.9 | 0.1 | 2.0 | 0.2 | 2.0 | 0.2 |
| Sodium | mEq/L | Saline | 144 | 3 | 143 | 1 | 144 | 1 | 142 | 2 | 144 | 1 | 141 | 2 | 144 | 2 |
|  |  | IGF-1 | 143 | 2 | 141 | 4 | 143 | 1 | 143 | 3 | 144 | 2 | 141 | 3 | 144 | 1 |
| Potassium | mEq/L | Saline | 4.0 | 0.3 | 3.9 | 0.1 | 4.4 | 0.5 | 4.2 | 0.2 | 4.1 | 0.3 | 4.2 | 0.2 | 4.6 | 0.1 |
|  |  | IGF-1 | 4.0 | 0.5 | 4.1 | 0.3 | 3.9 | 0.1 | 4.0 | 0.3 | 4.1 | 0.3 | 4.3 | 0.8 | 4.6 | 0.5 |
| NA/K RATIO |  | Saline | 36 | 3.5 | 36 | 1.3 | 33 | 3.4 | 34 | 2.1 | 35 | 2.1 | 34 | 1.5 | 32 | 1.3 |
|  |  | IGF-1 | 36 | 3.6 | 34 | 2.2 | 36 | 1.5 | 36 | 2.4 | 35 | 2.1 | 33 | 5.3 | 32 | 2.8 |
| Chloride | mEq/L | Saline | 99 | 4 | 99 | 2 | 99 | 1 | 100 | 1 | 103 | 2 | 101 | 2 | 102 | 1 |
|  |  | IGF-1 | 97 | 3 | 100 | 3 | 100 | 2 | 100 | 2 | 101 | 2 | 100 | 3 | 101 | 2 |
| Cholesterol | mg/dL | Saline | 150 | 42 | 361 | 83 | 366 | 62 | 331 | 52 | 307 | 48 | 289 | 131 | 250 | 56 |
|  |  | IGF-1 | 163 | 51 | 415 | 106 | 410 | 111 | 347 | 94 | 305 | 107 | 344 | 128 | 282 | 102 |
| TRIGLYCERIDE | mg/dL | Saline | 32 | 8 | 28 | 4 | 61 | 20 | 42 | 12 | 40 | 6 | 39 | 19 | 47 | 17 |
|  |  | IGF-1 | 33 | 14 | 37 | 23 | 29 | 10 | 30 | 5 | 32 | 5 | 34 | 8 | 41 | 7 |
| Amylase | IU/L | Saline | 1418 | 360 | 1603 | 360 | 1559 | 350 | 1441 | 343 | 1593 | 399 | 1546 | 411 | 1523 | 372 |
|  |  | IGF-1 | 1374 | 312 | 1459 | 340 | 1429 | 350 | 1381 | 379 | 1478 | 446 | 1501 | 416 | 1399 | 376 |
| PrecisionPSL | U/L | Saline | 7 | 1 | 7 | 1 | 8 | 2 | 6 | 1 | 6 | 2 | 6 | 1 | 7 | 1 |
|  |  | IGF-1 | 8 | 1 | 9 | 5 | 7 | 0 | 6 | 1 | 6 | 0 | 6 | 1 | 6 | 1 |
| CPK | IU/L | Saline | 408 | 121 | 2154 | 2160 | 3519 | 6915 | 498 | 112 | 621 | 359 | 753 | 914 | 381 | 54 |
|  |  | IGF-1 | 798 | 649 | 701 | 673 | 616 | 263 | 538 | 299 | 573 | 358 | 345 | 100 | 1715 | 3065 |

| Female | Units | Group | 0 |  | 1-mo |  | 2-mo |  | 3-mo |  | 4-mo |  | 5-mo |  | 6-mo |  |
|---|---|---|---|---|---|---|---|---|---|---|---|---|---|---|---|---|
|  |  |  | Avg | SD | Avg | SD | Avg | SD | Avg | SD | Avg | SD | Avg | SD | Avg | SD |
| Total Protein | g/dL | Saline | 6.6 | 0.5 | 6.7 | 0.5 | 6.6 | 0.4 | 6.6 | 0.3 | 6.9 | 0.5 | 7.0 | 0.6 | 6.8 | 0.3 |
|  |  | IGF-1 | 6.5 | 0.3 | 6.5 | 0.4 | 6.2 | 0.5 | 6.1 | 0.3 | 6.5 | 0.2 | 6.6 | 0.8 | 6.3 | 0.6 |
| Albumin | g/dL | Saline | 3.8 | 0.5 | 3.7 | 0.4 | 3.9 | 0.4 | 3.7 | 0.3 | 3.8 | 0.4 | 4.0 | 0.3 | 3.8 | 0.5 |
|  |  | IGF-1 | 3.8 | 0.2 | 3.6 | 0.3 | 3.9 | 0.2 | 3.7 | 0.3 | 4.1 | 0.3 | 3.8 | 0.6 | 3.6 | 0.4 |
| Globulin | g/dL | Saline | 2.8 | 0.3 | 2.9 | 0.6 | 2.6 | 0.5 | 2.9 | 0.5 | 3.1 | 0.6 | 3.1 | 0.6 | 3.0 | 0.4 |
|  |  | IGF-1 | 2.7 | 0.4 | 2.9 | 0.7 | 2.4 | 0.6 | 2.4 | 0.5 | 2.4 | 0.3 | 2.8 | 1.4 | 2.7 | 0.9 |
| A/G Ratio |  | Saline | 1.4 | 0.3 | 1.3 | 0.3 | 1.5 | 0.3 | 1.3 | 0.2 | 1.3 | 0.3 | 1.3 | 0.3 | 1.3 | 0.4 |
|  |  | IGF-1 | 1.5 | 0.4 | 1.3 | 0.4 | 1.7 | 0.5 | 1.6 | 0.5 | 1.7 | 0.3 | 1.6 | 0.7 | 1.5 | 0.5 |
| AST (SGOT) | IU/L | Saline | 18 | 4.3 | 20 | 2.4 | 20 | 1.5 | 20 | 2.1 | 20 | 3.8 | 28 | 12.3 | 20 | 7.0 |
|  |  | IGF-1 | 20 | 2.9 | 20 | 3.4 | 25 | 11.7 | 20 | 2.7 | 38 | 30.1 | 20 | 2.6 | 19 | 2.3 |
| ALT (SGPT) | IU/L | Saline | 25 | 3.0 | 27 | 3.4 | 27 | 3.4 | 23 | 3.3 | 21 | 1.3 | 21 | 2.3 | 21 | 3.2 |
|  |  | IGF-1 | 23 | 3.4 | 25 | 1.8 | 25 | 1.9 | 22 | 2.3 | 24 | 4.4 | 21 | 2.9 | 21 | 2.7 |
| Alk Phosphatase | IU/L | Saline | 65 | 15 | 75 | 23 | 69 | 19 | 64 | 21 | 60 | 23 | 67 | 18 | 61 | 15 |
|  |  | IGF-1 | 68 | 19 | 79 | 18 | 73 | 13 | 74 | 19 | 69 | 16 | 64 | 16 | 59 | 13 |
| GGT | IU/L | Saline | 25 | 6 | 26 | 9 | 24 | 9 | 37 | 13 | 25 | 9 | 28 | 8 | 28 | 7 |
|  |  | IGF-1 | 26 | 12 | 24 | 8 | 21 | 10 | 34 | 24 | 27 | 12 | 23 | 9 | 23 | 10 |
| Total Bilirubin | mg/dL | Saline | 0.1 | 0.1 | 0.1 | 0.0 | 0.1 | 0.0 | 0.1 | 0.1 | 0.1 | 0.0 | 0.1 | 0.0 | 0.1 | 0.0 |
|  |  | IGF-1 | 0.2 | 0.1 | 0.1 | 0.1 | 0.1 | 0.0 | 0.1 | 0.1 | 0.1 | 0.0 | 0.1 | 0.0 | 0.1 | 0.0 |
| BUN | mg/dL | Saline | 11 | 4.8 | 11 | 1.5 | 12 | 1.8 | 12 | 3.1 | 9 | 1.9 | 10 | 1.7 | 12 | 2.3 |
|  |  | IGF-1 | 10 | 1.9 | 8 | 1.5 | 7 | 0.7 | 7 | 1.1 | 8 | 1.8 | 7 | 1.5 | 8 | 1.5 |
| Creatinine | mg/dL | Saline | 1.5 | 0.2 | 1.6 | 0.1 | 1.5 | 0.2 | 1.7 | 0.2 | 1.8 | 0.2 | 1.8 | 0.2 | 1.8 | 0.3 |
|  |  | IGF-1 | 1.5 | 0.2 | 1.3 | 0.2 | 1.4 | 0.2 | 1.5 | 0.2 | 1.6 | 0.2 | 1.5 | 0.2 | 1.5 | 0.2 |
| BUN/CREAT RATIO |  | Saline | 8 | 3.9 | 7 | 1.0 | 8 | 1.9 | 7 | 1.9 | 5 | 1.7 | 6 | 1.3 | 7 | 2.1 |
|  |  | IGF-1 | 7 | 2.1 | 6 | 1.1 | 5 | 0.8 | 5 | 0.4 | 5 | 1.3 | 5 | 0.5 | 5 | 0.8 |
| Phosphorus | mg/dL | Saline | 5.8 | 0.3 | 5.7 | 0.2 | 5.8 | 0.2 | 5.8 | 0.8 | 5.6 | 0.4 | 5.6 | 0.4 | 5.8 | 0.7 |
|  |  | IGF-1 | 5.8 | 0.1 | 6.4 | 0.5 | 5.7 | 0.3 | 5.7 | 0.6 | 5.9 | 0.5 | 5.5 | 0.6 | 5.4 | 0.5 |
| Glucose | mg/dL | Saline | 141 | 9 | 112 | 24 | 101 | 14 | 146 | 54 | 112 | 33 | 91 | 10 | 113 | 13 |
|  |  | IGF-1 | 128 | 23 | 93 | 11 | 100 | 21 | 104 | 29 | 100 | 12 | 84 | 13 | 86 | 11 |
| CALCIUM | mg/dL | Saline | 10.8 | 0.5 | 9.9 | 0.3 | 10.4 | 0.4 | 10.4 | 0.2 | 10.4 | 0.3 | 10.5 | 0.2 | 10.4 | 0.3 |
|  |  | IGF-1 | 10.7 | 0.2 | 9.9 | 0.4 | 10.2 | 0.2 | 10.3 | 0.1 | 10.3 | 0.2 | 10.1 | 0.3 | 9.8 | 0.4 |
| Magnesium | mEq/L | Saline | 2.0 | 0.2 | 2.0 | 0.3 | 2.1 | 0.3 | 1.9 | 0.3 | 1.9 | 0.2 | 1.9 | 0.2 | 2.0 | 0.1 |
|  |  | IGF-1 | 1.8 | 0.1 | 1.8 | 0.2 | 1.8 | 0.1 | 1.7 | 0.1 | 1.8 | 0.2 | 1.6 | 0.1 | 1.6 | 0.1 |
| Sodium | mEq/L | Saline | 141 | 3 | 140 | 2 | 138 | 3 | 135 | 4 | 139 | 3 | 138 | 2 | 139 | 4 |
|  |  | IGF-1 | 140 | 1 | 140 | 2 | 139 | 0 | 140 | 2 | 141 | 2 | 140 | 1 | 139 | 2 |
| Potassium | mEq/L | Saline | 4.0 | 0.1 | 4.0 | 0.2 | 4.1 | 0.6 | 4.2 | 0.4 | 4.3 | 0.6 | 3.9 | 0.2 | 4.2 | 0.5 |
|  |  | IGF-1 | 3.9 | 0.1 | 4.3 | 0.3 | 4.0 | 0.3 | 4.0 | 0.1 | 4.7 | 0.9 | 4.2 | 0.3 | 4.1 | 0.2 |
| NA/K RATIO |  | Saline | 35 | 1.2 | 35 | 2.5 | 34 | 4.1 | 32 | 2.9 | 33 | 4.1 | 36 | 1.5 | 34 | 3.8 |
|  |  | IGF-1 | 36 | 1.3 | 33 | 2.5 | 35 | 2.5 | 35 | 0.8 | 31 | 5.4 | 34 | 2.2 | 34 | 1.9 |
| Chloride | mEq/L | Saline | 99 | 3 | 100 | 2 | 97 | 3 | 94 | 3 | 99 | 3 | 98 | 3 | 98 | 3 |
|  |  | IGF-1 | 99 | 2 | 99 | 2 | 100 | 1 | 101 | 1 | 101 | 2 | 101 | 1 | 102 | 1 |
| Cholesterol | mg/dL | Saline | 441 | 153 | 932 | 337 | 917 | 359 | 922 | 428 | 812 | 329 | 860 | 367 | 809 | 293 |
|  |  | IGF-1 | 427 | 95 | 696 | 96 | 764 | 196 | 764 | 173 | 723 | 181 | 655 | 220 | 615 | 139 |
| TRIGLYCERIDE | mg/dL | Saline | 83 | 26 | 54 | 13 | 61 | 10 | 61 | 10 | 57 | 17 | 63 | 25 | 91 | 31 |
|  |  | IGF-1 | 55 | 22 | 48 | 35 | 45 | 11 | 40 | 14 | 49 | 19 | 48 | 35 | 38 | 8 |
| Amylase | IU/L | Saline | 1199 | 267 | 1188 | 479 | 1192 | 464 | 1154 | 420 | 1128 | 387 | 1284 | 546 | 1271 | 475 |
|  |  | IGF-1 | 1283 | 502 | 1313 | 467 | 1355 | 514 | 1287 | 504 | 1283 | 448 | 1340 | 537 | 1378 | 561 |
| PrecisionPSL | U/L | Saline | 10 | 5 | 8 | 1 | 8 | 2 | 8 | 1 | 8 | 1 | 8 | 2 | 9 | 1 |
|  |  | IGF-1 | 8 | 0 | 8 | 2 | 7 | 0 | 7 | 0 | 9 | 3 | 7 | 1 | 9 | 1 |
| CPK | IU/L | Saline | 260 | 74 | 286 | 33 | 343 | 95 | 338 | 90 | 408 | 345 | 919 | 985 | 282 | 61 |
|  |  | IGF-1 | 509 | 412 | 347 | 155 | 874 | 1245 | 330 | 74 | 1405 | 2095 | 603 | 818 | 271 | 91 |

**Supplemental Table 3A. FH pigs CBC with differential (males)**

| Male Complete Blood Count | Units | Group | 0 Avg | 0 SD | 1-mo Avg | 1-mo SD | 2-mo Avg | 2-mo SD | 3-mo Avg | 3-mo SD | 4-mo Avg | 4-mo SD | 5-mo Avg | 5-mo SD | 6-mo Avg | 6-mo SD |
|---|---|---|---|---|---|---|---|---|---|---|---|---|---|---|---|---|
| WBC | 10^3/uL | Saline | 9.2 | 2.4 | 6.7 | 1.0 | 7.4 | 0.7 | 7.4 | 1.3 | 8.0 | 2.0 | 7.9 | 1.4 | 8.6 | 1.6 |
| | | IGF-1 | 9.1 | 2.8 | 8.0 | 1.2 | 7.4 | 0.7 | 8.9 | 1.1 | 7.9 | 1.0 | 7.7 | 1.5 | 8.6 | 0.9 |
| RBC | 10^6/uL | Saline | 6.0 | 1.6 | 6.1 | 0.7 | 5.6 | 0.5 | 4.8 | 0.5 | 5.1 | 0.5 | 4.9 | 0.4 | 4.7 | 0.5 |
| | | IGF-1 | 5.3 | 0.8 | 6.1 | 0.9 | 5.0 | 0.7 | 4.4 | 0.5 | 4.9 | 0.5 | 4.6 | 0.4 | 4.2 | 0.6 |
| HGB | g/dL | Saline | 11.4 | 2.7 | 12.0 | 1.3 | 11.2 | 0.9 | 10.0 | 1.1 | 10.5 | 1.1 | 10.4 | 0.8 | 9.8 | 1.0 |
| | | IGF-1 | 10.2 | 1.5 | 11.9 | 1.5 | 10.1 | 1.0 | 8.9 | 0.7 | 10.0 | 0.7 | 9.9 | 0.9 | 8.9 | 1.3 |
| HCT | % | Saline | 38 | 9 | 38 | 4 | 37 | 3 | 30 | 3 | 34 | 3 | 33 | 2 | 31 | 3 |
| | | IGF-1 | 33 | 5 | 38 | 5 | 31 | 4 | 27 | 3 | 33 | 3 | 31 | 3 | 29 | 4 |
| MCV | fL | Saline | 64 | 3 | 63 | 1 | 66 | 2 | 63 | 6 | 66 | 2 | 67 | 1 | 65 | 3 |
| | | IGF-1 | 63 | 4 | 62 | 2 | 63 | 2 | 62 | 6 | 67 | 1 | 67 | 2 | 68 | 2 |
| MCH | pg | Saline | 19.2 | 0.8 | 19.7 | 0.7 | 20.0 | 0.8 | 20.8 | 0.7 | 20.5 | 0.7 | 21.2 | 0.6 | 20.8 | 0.9 |
| | | IGF-1 | 19.1 | 0.8 | 19.7 | 0.9 | 20.7 | 1.1 | 20.2 | 1.6 | 20.5 | 0.7 | 21.5 | 1.0 | 21.3 | 0.9 |
| MCHC | g/dL | Saline | 30 | 0.9 | 31 | 0.4 | 30 | 0.5 | 33 | 2.4 | 31 | 0.4 | 32 | 1.1 | 32 | 2.1 |
| | | IGF-1 | 30 | 0.7 | 32 | 0.5 | 33 | 1.4 | 33 | 4.1 | 30 | 0.5 | 32 | 0.8 | 31 | 0.8 |
| Platelet Count | 10^3/uL | Saline | 382 | 66 | 326 | 48 | 371 | 31 | 340 | 65 | 301 | 23 | 349 | 55 | 386 | 74 |
| | | IGF-1 | 346 | 65 | 253 | 41 | 274 | 104 | 258 | 86 | 324 | 74 | 289 | 43 | 305 | 79 |
| Differential (absolute) | Units | | | | | | | | | | | | | | | |
| Neutrophils | /uL | Saline | 3790 | 2260 | 2105 | 780 | 2608 | 751 | 2750 | 1045 | 3346 | 1457 | 3239 | 1126 | 3183 | 1680 |
| | | IGF-1 | 4448 | 2162 | 3163 | 1291 | 2616 | 535 | 3594 | 308 | 3700 | 1259 | 3393 | 1165 | 3245 | 661 |
| Bands | | | | | | | | | | | | | | | | |
| Lymphocytes | /uL | Saline | 5017 | 757 | 4306 | 350 | 4518 | 559 | 4189 | 536 | 4239 | 875 | 4267 | 644 | 5140 | 1062 |
| | | IGF-1 | 4261 | 1521 | 4366 | 787 | 4516 | 774 | 4767 | 834 | 3618 | 640 | 3967 | 503 | 4700 | 127 |
| Monocytes | /uL | Saline | 352 | 109 | 254 | 62 | 284 | 50 | 321 | 114 | 415 | 265 | 295 | 61 | 170 | 151 |
| | | IGF-1 | 321 | 82 | 355 | 252 | 244 | 110 | 383 | 176 | 541 | 228 | 307 | 59 | 293 | 120 |
| Eosinophils | /uL | Saline | 50 | 47 | 55 | 32 | 30 | 41 | 120 | 51 | 39 | 54 | 99 | 66 | 106 | 56 |
| | | IGF-1 | 110 | 53 | 77 | 59 | 64 | 71 | 156 | 64 | 80 | 96 | 0 | 0 | 322 | 346 |
| Basophils | /uL | Saline | 0 | 0 | 0 | 0 | 0 | 0 | 0 | 0 | 0 | 0 | 0 | 0 | 0 | 0 |
| | | IGF-1 | 0 | 0 | 0 | 0 | 0 | 0 | 0 | 0 | 0 | 0 | 12 | 28 | 0 | 0 |

**Supplemental Table 3B. FH pigs CBC with differential (females)**

| Female | | | 0 | | 1-mo | | 2-mo | | 3-mo | | 4-mo | | 5-mo | | 6-mo | |
|---|---|---|---|---|---|---|---|---|---|---|---|---|---|---|---|---|
| **Complete Blood Count** | **Units** | **Group** | **Avg** | **SD** | **Avg** | **SD** | **Avg** | **SD** | **Avg** | **SD** | **Avg** | **SD** | **Avg** | **SD** | **Avg** | **SD** |
| WBC | 10^3/uL | Saline | 7.1 | 3.3 | 6.2 | 1.8 | 6.9 | 2.5 | 6.4 | 2.0 | 7.1 | 3.4 | 7.6 | 2.9 | 6.7 | 3.0 |
| | | IGF-1 | 5.9 | 0.6 | 6.8 | 0.9 | 6.4 | 1.2 | 6.3 | 1.3 | 7.2 | 0.8 | 7.7 | 1.9 | 6.9 | 1.9 |
| RBC | 10^6/uL | Saline | 4.4 | 0.4 | 5.0 | 0.7 | 4.8 | 0.6 | 4.6 | 0.4 | 4.7 | 0.7 | 5.3 | 0.9 | 4.6 | 0.6 |
| | | IGF-1 | 4.8 | 0.3 | 4.8 | 0.4 | 5.1 | 0.7 | 4.6 | 0.3 | 4.9 | 0.5 | 4.6 | 0.7 | 4.3 | 0.2 |
| HGB | g/dL | Saline | 8.4 | 0.4 | 9.3 | 1.0 | 9.2 | 0.8 | 8.9 | 0.8 | 9.2 | 1.3 | 10.3 | 1.3 | 8.9 | 0.9 |
| | | IGF-1 | 9.1 | 0.6 | 9.3 | 0.9 | 10.0 | 1.4 | 9.2 | 0.6 | 9.9 | 1.2 | 9.3 | 1.3 | 8.7 | 0.4 |
| HCT | % | Saline | 26 | 2 | 29 | 3 | 28 | 3 | 26 | 2 | 28 | 4 | 31 | 4 | 27 | 3 |
| | | IGF-1 | 28 | 3 | 29 | 3 | 30 | 4 | 27 | 2 | 30 | 3 | 28 | 4 | 26 | 2 |
| MCV | fL | Saline | 59 | 4 | 58 | 2 | 58 | 3 | 57 | 3 | 59 | 3 | 59 | 3 | 60 | 4 |
| | | IGF-1 | 59 | 4 | 60 | 3 | 59 | 1 | 59 | 1 | 61 | 1 | 61 | 3 | 62 | 1 |
| MCH | pg | Saline | 19.0 | 1.0 | 18.7 | 1.3 | 19.5 | 1.3 | 21.5 | 5.1 | 19.4 | 1.1 | 19.4 | 1.0 | 19.7 | 1.5 |
| | | IGF-1 | 18.9 | 0.5 | 19.5 | 0.7 | 19.6 | 0.8 | 20.1 | 0.5 | 20.4 | 0.5 | 20.2 | 0.5 | 20.1 | 0.4 |
| MCHC | g/dL | Saline | 33 | 2.6 | 32 | 1.6 | 34 | 0.8 | 34 | 0.8 | 33 | 0.4 | 33 | 0.4 | 33 | 0.8 |
| | | IGF-1 | 32 | 2.1 | 33 | 0.8 | 33 | 1.4 | 34 | 0.5 | 33 | 0.4 | 34 | 1.0 | 33 | 0.9 |
| Platelet Count | 10^3/uL | Saline | 325 | 37 | 347 | 62 | 320 | 87 | 368 | 83 | 375 | 143 | 327 | 68 | 318 | 60 |
| | | IGF-1 | 290 | 110 | 345 | 112 | 280 | 99 | 286 | 95 | 257 | 139 | 243 | 172 | 282 | 122 |
| Differential (absolute) | Units | | | | | | | | | | | | | | | |
| Neutrophils | /uL | Saline | 2904 | 2995 | 2366 | 1977 | 2477 | 2385 | 2454 | 2213 | 2802 | 2908 | 2906 | 2846 | 2509 | 2457 |
| | | IGF-1 | 1744 | 513 | 2489 | 1232 | 1535 | 1034 | 1455 | 900 | 1879 | 387 | 2466 | 1618 | 2234 | 1589 |
| Bands | | | | | | | | | | | | | | | | |
| Lymphocytes | /uL | Saline | 3985 | 1674 | 3666 | 1076 | 4174 | 645 | 3597 | 774 | 3930 | 809 | 4148 | 846 | 3740 | 520 |
| | | IGF-1 | 3683 | 778 | 3845 | 715 | 4455 | 1097 | 4506 | 754 | 4869 | 989 | 4737 | 1014 | 4407 | 467 |
| Monocytes | /uL | Saline | 147 | 40 | 167 | 89 | 185 | 129 | 272 | 80 | 270 | 93 | 367 | 99 | 292 | 123 |
| | | IGF-1 | 319 | 205 | 384 | 167 | 416 | 162 | 247 | 154 | 234 | 81 | 361 | 218 | 182 | 68 |
| Eosinophils | /uL | Saline | 84 | 80 | 0 | 0 | 24 | 54 | 56 | 61 | 79 | 52 | 139 | 109 | 91 | 42 |
| | | IGF-1 | 109 | 84 | 41 | 93 | 14 | 31 | 77 | 39 | 178 | 164 | 136 | 144 | 76 | 67 |
| Basophils | /uL | Saline | 0 | 0 | 0 | 0 | 0 | 0 | 0 | 0 | 0 | 0 | 0 | 0 | 47 | 106 |
| | | IGF-1 | 25 | 35 | 0 | 0 | 0 | 0 | 14 | 32 | 0 | 0 | 0 | 0 | 0 | 0 |

**Supplemental Figure 2A**

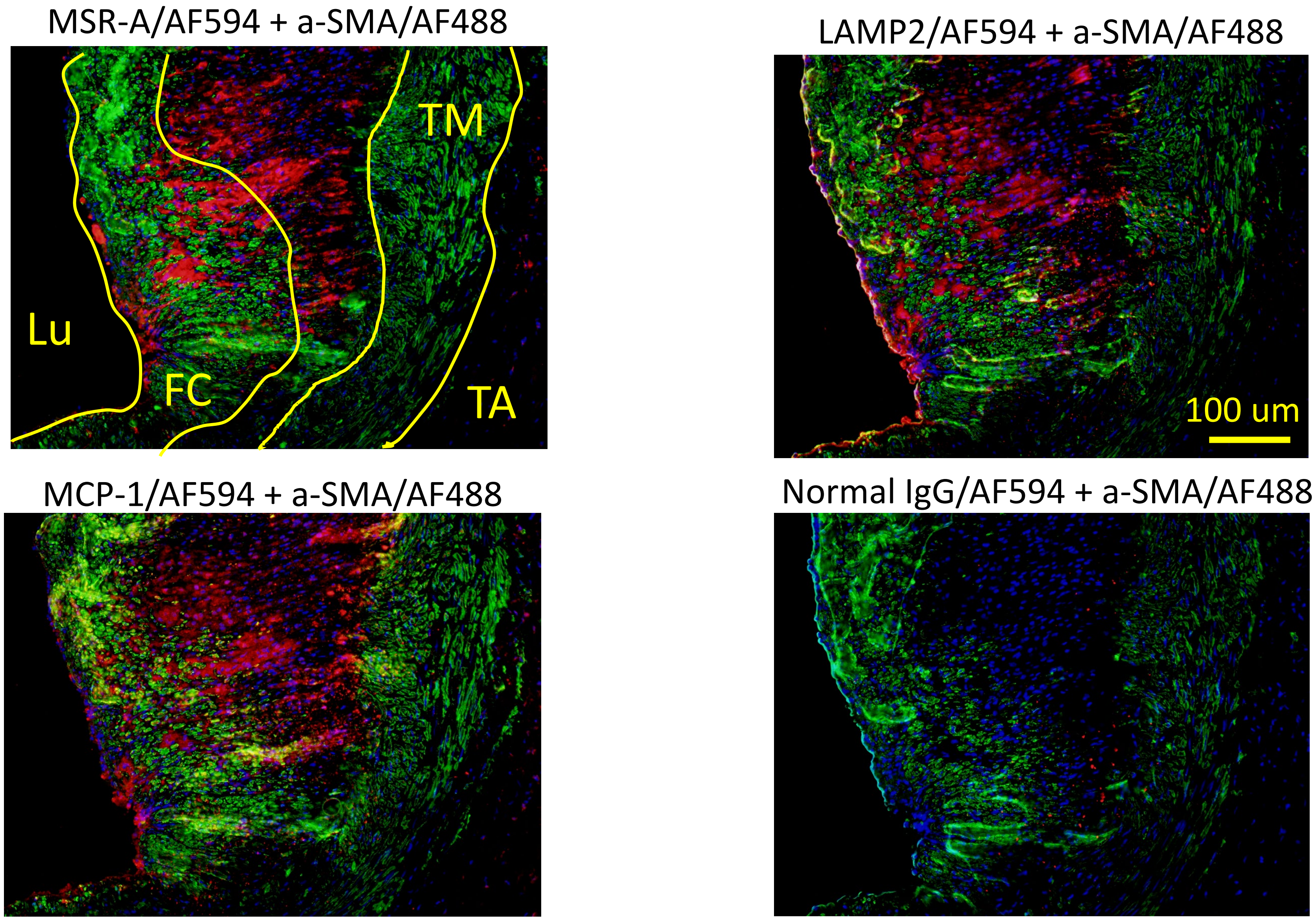

**Supplemental Figure 2B**

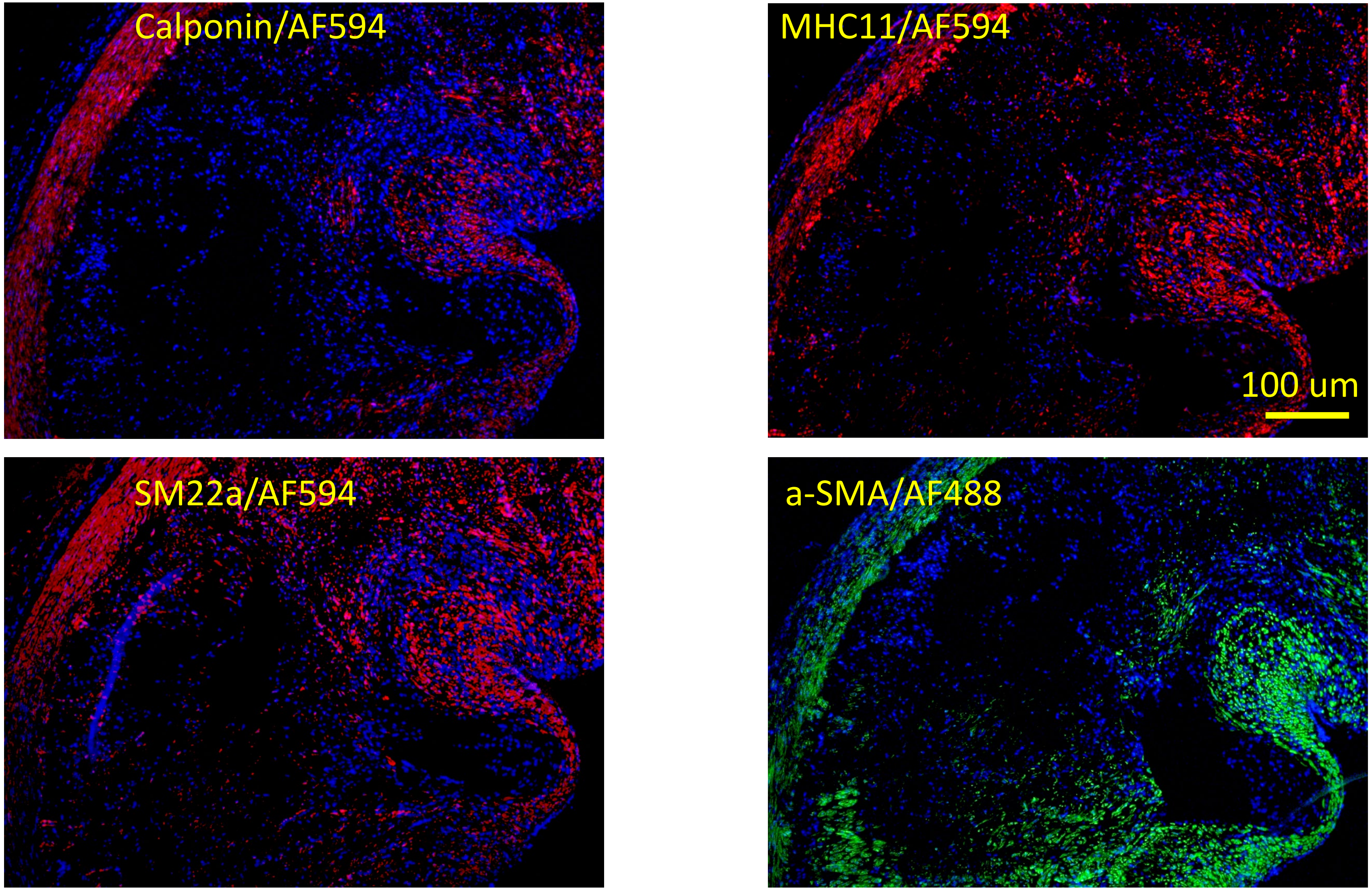

**Supplemental Figure 2C**

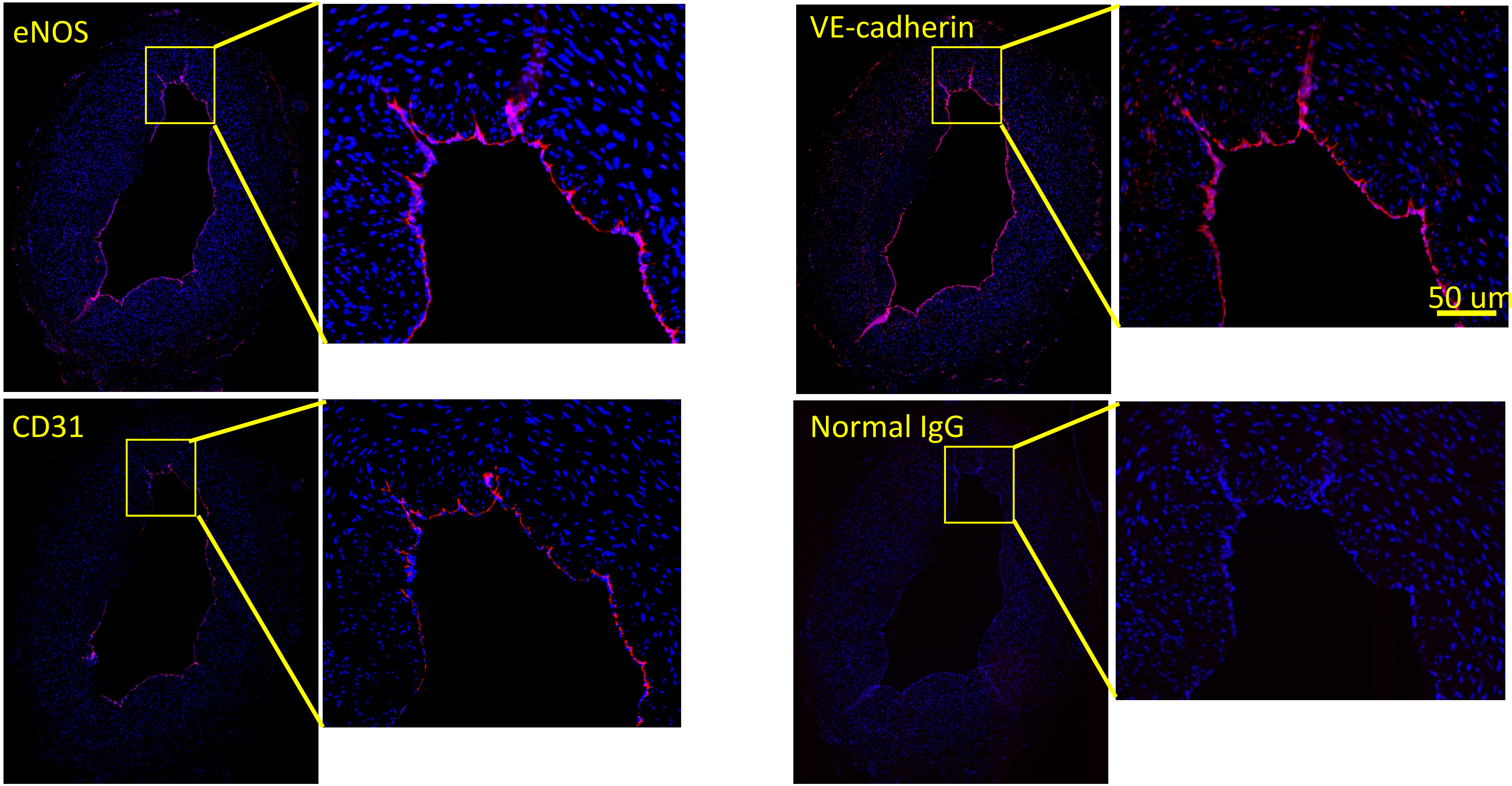

**Supplemental Figure 3**

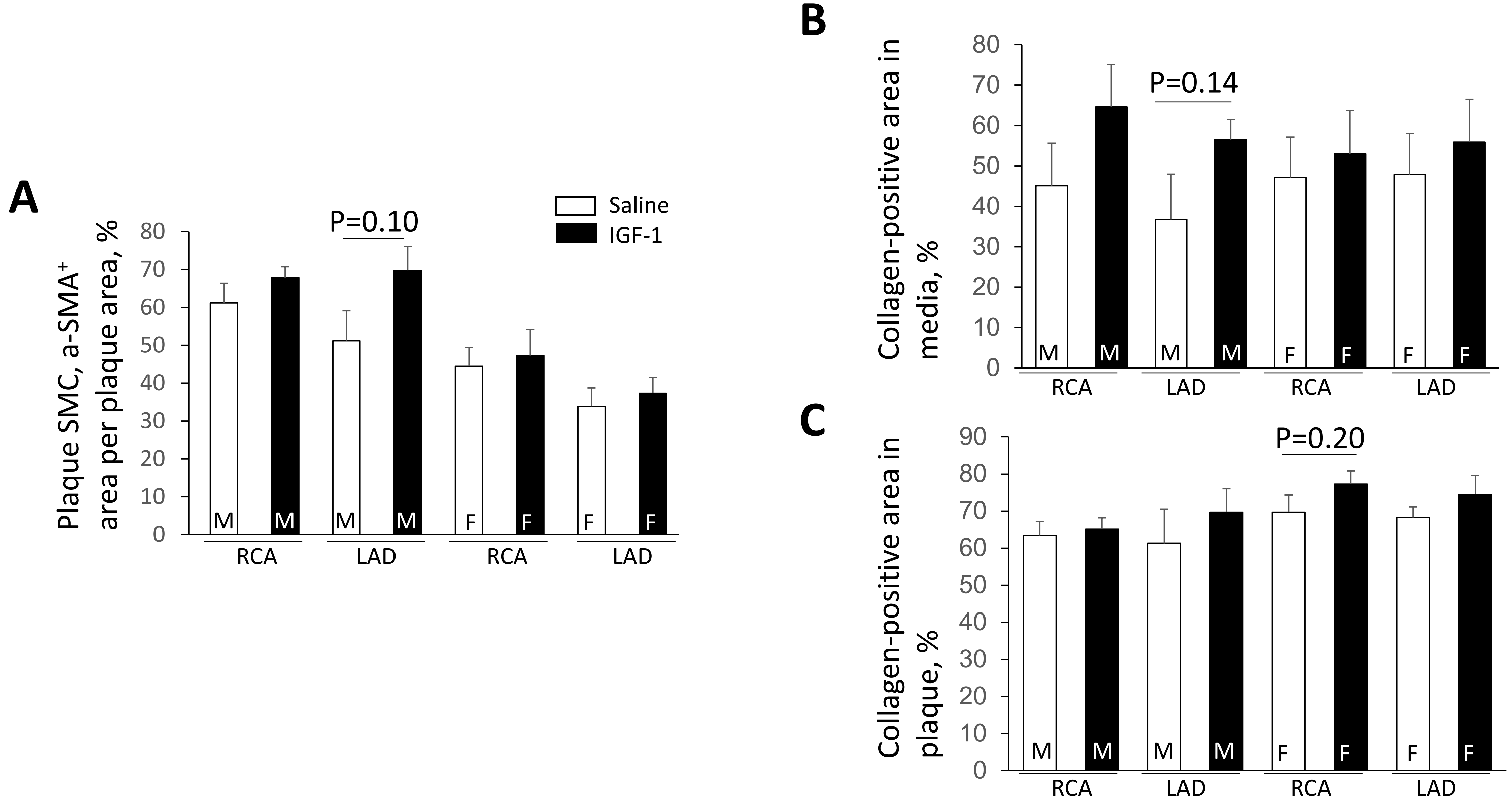

#### Supplemental Table 4. Spatial transcriptomic quality controls

| ST quality control | Mean | SEM |
|---|---|---|
| Number of Reads, millions | 121.0 | 14.6 |
| Median Genes per Spot | 2122.4 | 131.4 |
| Median UMI Counts per Spot | 6477.5 | 652.9 |
| Valid Barcodes, % | 98.0 | 0.1 |
| Valid UMIs, % | 99.9 | 0.001 |
| Q30 Bases in Barcode, % | 96.0 | 0.1 |
| Q30 Bases in RNA Read, % | 95.6 | 0.001 |
| Q30 Bases in UMI, % | 96.5 | 0.1 |
| Reads Mapped to Genome, % | 94.2 | 0.9 |
| Reads Mapped Confidently to Genome, % | 91.7 | 1.0 |
| Reads Mapped Confidently to Intergenic Regions, % | 3.0 | 0.2 |
| Reads Mapped Confidently to Intronic Regions, % | 5.7 | 0.5 |
| Reads Mapped Confidently to Exonic Regions, % | 83.0 | 1.6 |
| Reads Mapped Confidently to Transcriptome, % | 77.2 | 1.7 |
| Reads Mapped Antisense to Gene, % | 1.2 | 0.1 |
| Fraction Reads in Spots Under Tissue, % | 82.7 | 1.2 |
| Total Genes Detected | 17146 | 283 |

**Supplemental Figure 4**

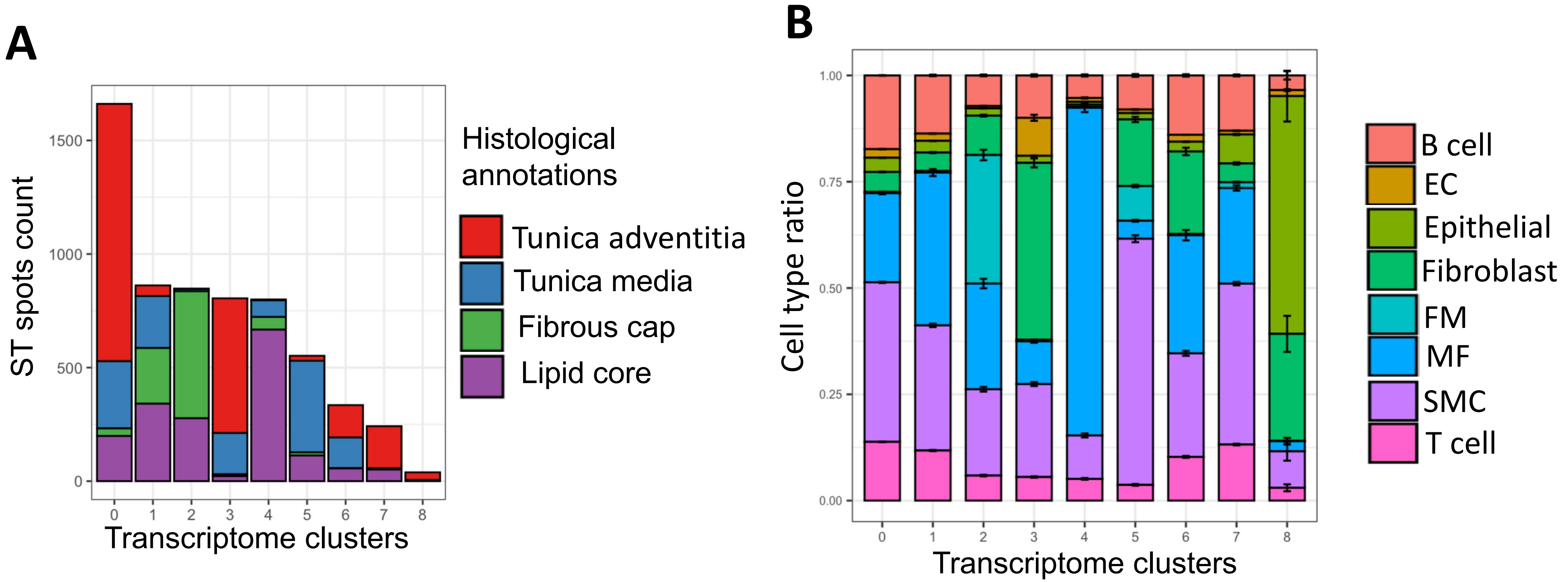
